# Transcriptomic Profiling of Diabetic Porcine Wound Healing Model Identifies Key Metabolic, Inflammatory, and Oxidative Stress Pathways

**DOI:** 10.64898/2026.01.30.702870

**Authors:** Joshua T. McCune, Mariah G. Bezold, Jeffrey M. Davidson, C. Henrique Serezani, Rebecca S. Cook, Craig L. Duvall

**Affiliations:** Department of Biomedical Engineering, Vanderbilt University, Nashville, TN, USA; Department of Pathology, Microbiology and Immunology, Vanderbilt University School of Medicine, Nashville, TN, USA; Department of Medicine, Division of Infectious Diseases, Vanderbilt University Medical Center, Nashville

**Keywords:** diabetic foot ulcer, porcine model, streptozotocin, RNA-seq, wound healing

## Abstract

Diabetic foot ulcers (**DFU**s) remain a major clinical challenge as diabetes prevalence rises, emphasizing the need for improved therapeutics and relevant preclinical models. Common rodent wound-healing models poorly recapitulate human skin anatomy and repair. Although porcine skin is comparable to human skin, many studies employ young, healthy pigs that do not reflect typical chronic human wounds. Here, we evaluated wound healing in full-thickness skin wounds in non-diabetic and diabetic Yucatan minipigs. RNA sequencing identified key transcriptional differences in wounds of diabetic versus non-diabetic animals, including pathways linked to increased inflammation and oxidative stress, as well as decreased metabolism and extracellular matrix organization, known hallmarks of DFUs. These findings support this preclinical model as a powerful approach for discovery and therapeutic testing in diabetic wounds and provide a novel data set for further mining of potential gene targets for diabetic wound intervention.

## Introduction

Chronic wounds, especially diabetic foot ulcers (**DFU**s), are associated with high rates of lower limb amputation and increased patient mortality^1^. The DFU microenvironment is complex – involving persistent inflammation, dysregulation of fibroblasts and keratinocytes, imbalanced extracellular matrix (**ECM**) production, oxidative stress, and metabolic dysfunction, all of which delay wound healing^2^. With an aging population and rising diabetes rates, there is a pressing need to improve DFU therapies. However, testing of new treatments and drugs is limited to a few suitable animal models that recapitulate the diabetic condition and molecular factors driving chronic DFUs. Although commonly used diabetic mouse wound-healing models (e.g., streptozotocin-induced, db/db, NOD) have provided valuable insight, they lack key features of human skin and heal primarily through contraction, necessitating stenting for accurate wound-healing studies^2^. Porcine wound-healing models are more relevant due to their anatomical and physiological similarities to human skin; however, comprehensive characterization of diabetic porcine models remains limited^2,3^. In this study, we have characterized a streptozotocin-induced (**STZ**) porcine model of full-thickness wound healing, detailing delayed wound closure and, for the first time, comparing bulk transcriptomic differences across diabetic and non-diabetic wounds within this model.

## Materials and Methods

All materials and methods are detailed in the Supporting Information Materials and Methods section.

## Results and Discussion

### Streptozotocin-induced diabetes impairs wound healing in Yucatan minipigs

We induced diabetes in a Yucatan minipig via intravenous infusion of STZ (125 mg/kg), followed by a 60-day maturation period with supplemental insulin to establish a hyperglycemic state while maintaining animal viability (**Figure 1A**). The diabetic state was maintained throughout the study, with blood glucose levels in the hyperglycemic range (>200 mg/dL) (**Figure 1B; Figure S1A-B**). Insulin was discontinued one week before wounding to mirror clinical scenarios in which diabetes remains poorly controlled and promote the underlying pathophysiology of chronic wounds. Full-thickness skin wounds were created 60 days after diabetes induction using a 2 cm biopsy punch. Identical wounds were generated in a non-diabetic Yucatan minipig to compare the effect of diabetes on wound healing in this model. Wound area was measured over 10 days, after which the study was terminated, capturing an intermediate wound-healing timepoint to assess differences in the resolution of inflammation, granulation tissue formation, and re-epithelialization. The diabetic wounds demonstrated significant delays in wound closure and re-epithelialization, quantified through digital planimetry and immunohistochemistry for cytokeratin 14, respectively, compared to non-diabetic wounds (**Figure 1C-E; Figure S1C**). Histological examination of diabetic wounds revealed striking differences in immune cell infiltration and phenotype with reduced collagen/granulation tissue deposition compared with non-diabetic wounds (**Figure 1F, Figure 2F; Figure S2**). These results highlight diabetes-associated wound-healing impairments, including delayed wound closure, diminished matrix deposition, and decreased re-epithelialization, closely resembling the wound-healing deficits that characterize human DFUs^4-6^.

**Figure 1.**
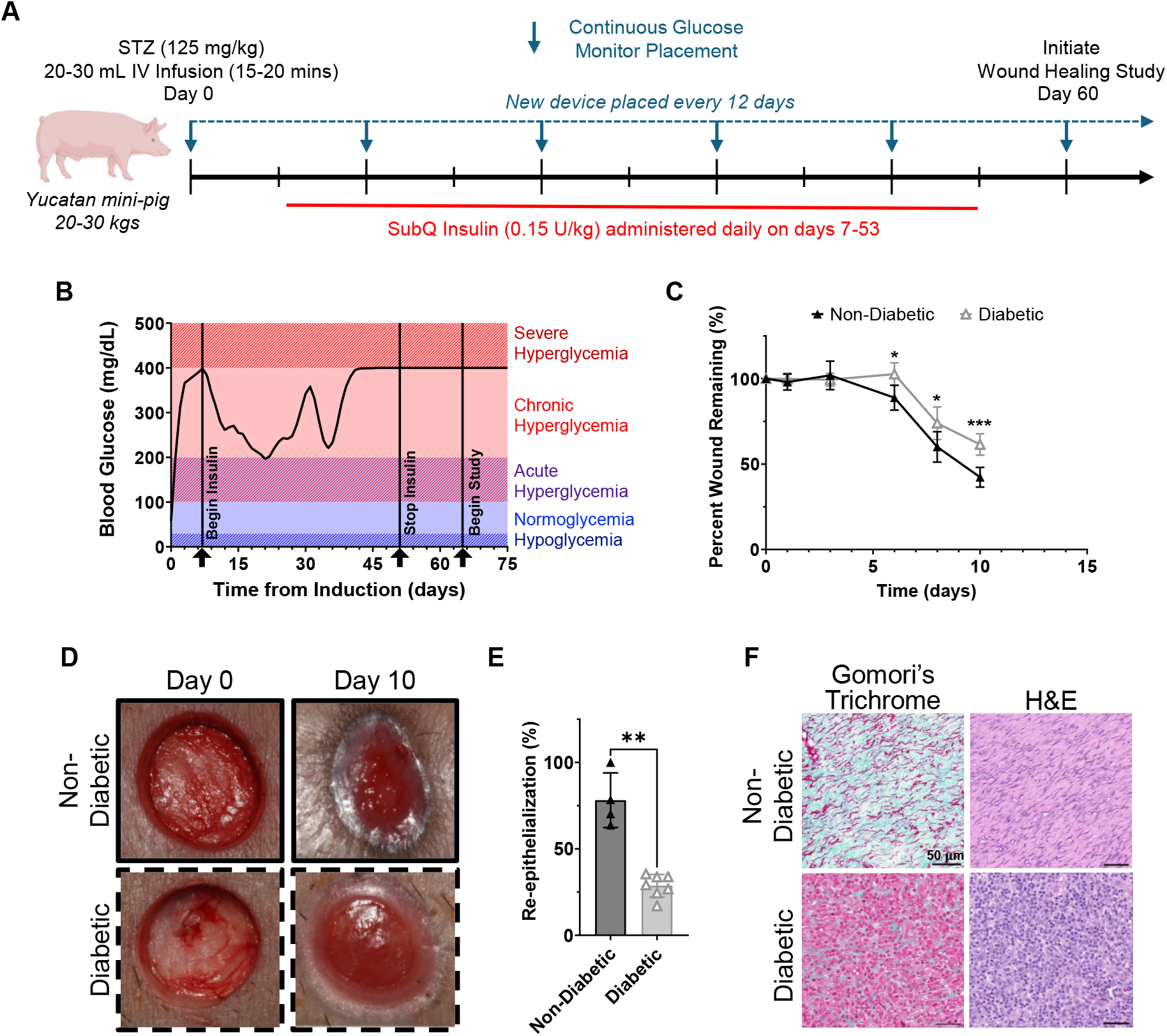
Streptozotocin induction and wound healing study in Yucatan minipig. **(A)** Scheme and timeline of STZ induction and wound healing study, **(B)** daily average blood glucose concentration of diabetic minipig, **(C)** full-thickness wound closure rates over time quantified from macroscopic photographs of the wounds, **(D)** representative images of wounds at day 0 and 10, **(E)** re-epithelialization of wounds at day 10 quantified from immunohistochemistry of cytokeratin14, and **(F)** representative histology of wounds at day 10. *P<0.05, **P<0.01, and ***P<0.001, by Welch’s unpaired t-test.

**Figure 2.**
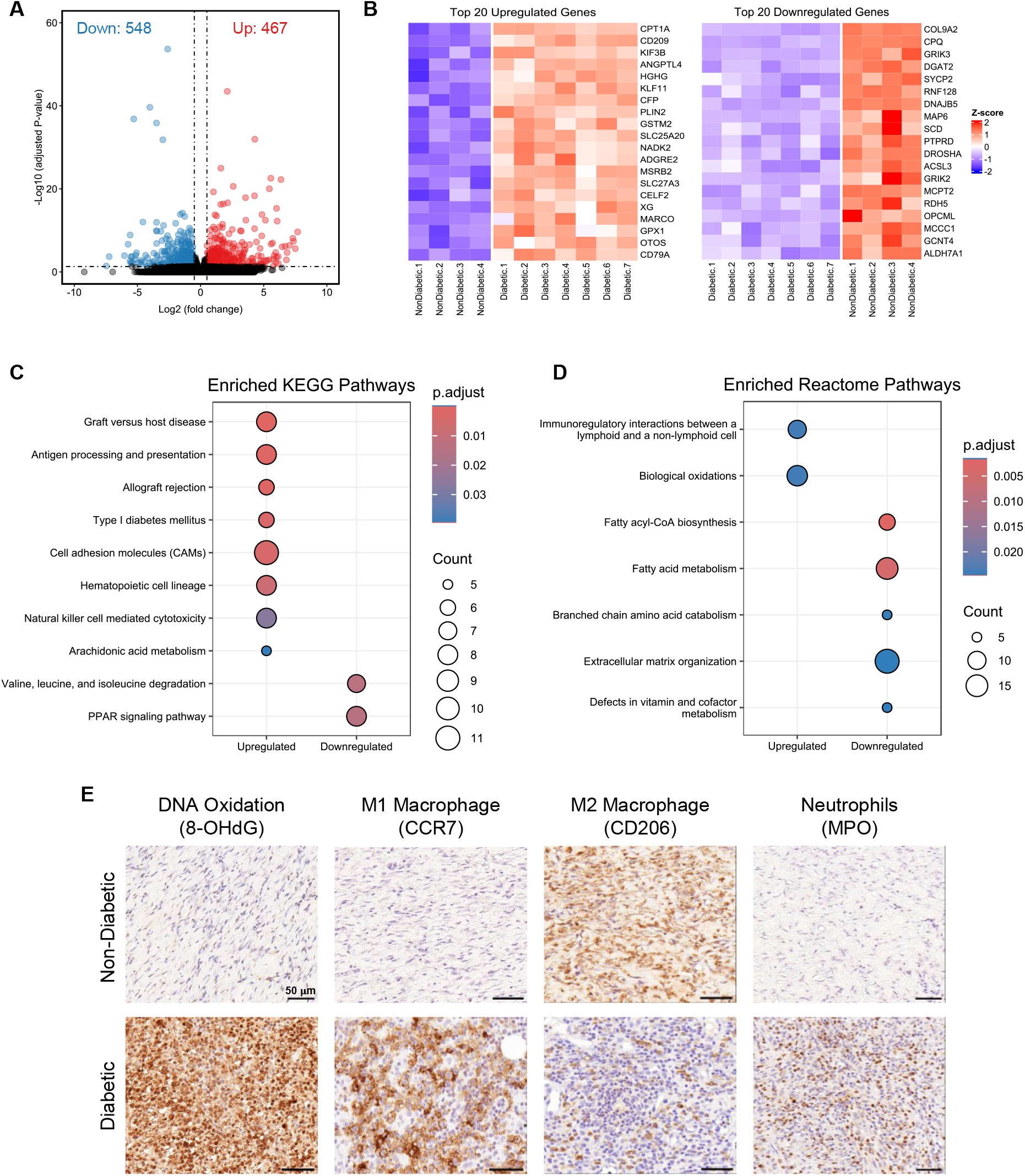
Differential gene expression analysis, functional enrichment analysis, and immunohistochemistry (IHC) comparing non-diabetic and diabetic wounds. **(A)** Volcano plot of differentially expressed genes, **(B)** heatmaps of top 20 upregulated and downregulated genes within the gene set, **(C)** KEGG enrichment analysis results, **(D)** Reactome enrichment analysis results, and **(E)** representative IHC of immune cell populations in diabetic and non-diabetic wounds at day 10.

### Wounds in diabetic minipigs exhibit different transcriptional profiles than non-diabetic wounds

Bulk RNA sequencing of diabetic wounds (N = 7) at day 10 revealed distinct gene expression profiles compared to non-diabetic wounds (N = 4), with 1,015 genes demonstrating differential expression (Log2FC > 0.5, adjusted p < 0.05) (**Figure 2A**). The 20 most activated and suppressed genes (**Figure 2B**) include those related to metabolic dysfunction (CPT1A, PLIN2, SLC25A20, NADK2, SLC27A3, CPQ, DGAT2, SCD, PTPRD, ACSL3, MCCC1, ALDH7A1), oxidative stress (GST, MSRB2, GPX1), and inflammation (CD209, IGHG, CFP, CELF2, MARCO, CD79A, RNF128). Functional enrichment analysis further corroborated these molecular signatures, revealing four major categories of dysregulated pathways (**Figure 2B-C; Figures S3-16**).

### Inflammation and immune dysregulation characterize wounds of diabetic minipig

Upregulated genes related to innate and adaptive immunity were enriched in autoimmune-associated KEGG pathways, consistent with STZ-induced diabetes (graft-versus-host disease, allograft rejection, and Type I diabetes mellitus)^7-9^. Critical genes enriched in these pathways include TNF and major histocompatibility (**MHC**) complex genes (SLA-DOB, SLA-DMA, SLA-8, SLA-DRA). This enrichment aligns with the mechanism of STZ-induced autoimmunity, which triggers MHC upregulation and the generation of auto-reactive lymphocytes^10^. These upregulated genes also implicate activated antigen presentation and immune surveillance within diabetic wounds, consistent with hyperglycemia-induced changes in macrophage-mediated immune responses^11,12^.

Genes encoding cell adhesion molecules (**CAM**s) involved in immune cell extravasation and infiltration into tissue (LTCAM, ITGAM, SELL, ICAM3) were significantly upregulated in diabetic wounds. Elevated levels of immune cell infiltration were also detected through increased CCR7 and myeloperoxidase (**MPO**) immunostaining, reflecting increased M1 pro-inflammatory macrophage and neutrophil infiltration, respectively, in diabetic wounds (**Figure 2E**). Similarly, hematopoietic cell lineage pathways were enriched in diabetic wounds, indicating local immune cell expansion (FCER2, CSF2RA, CD14, ITGAM, CD22, SLA-DRA, TNF, and IL1R2). Reactome enrichment analysis further suggested that diabetic wounds harbored excessive, pathological pro-inflammatory immune activation. Transcriptomic conclusions were corroborated by the elevated cellularity and immune infiltration observed histologically. The natural killer (**NK**) cell-mediated cytotoxicity pathway was also enriched. Strikingly, enriched genes represented both activating and inhibitory NK receptors (TNF, FCGR3A, SLA-8, PIK3R5, SHC2, KLRD1, RAC2, and KLRC1). These findings are consistent with observed NK cell dysregulation in diabetes, contributing to oxidative stress and inflammatory tissue damage^13,14^.

### Oxidative stress is upregulated in wounds of diabetic minipig

Enrichment in arachidonic acid metabolism and biological oxidation pathways, with upregulation of oxidative response genes (PTGES, PTGS1, CYP2B6B, GPX1, and GGT5) is consistent with elevated ROS burden in diabetic wounds. This hypothesis was confirmed by 8-OHdG immunostaining, reflecting increased DNA oxidative damage in diabetic wounds (**Figure 2E; Figure S2A**). Elevated ROS in diabetic wounds has been noted previously noted and is multifactorial, but hyperglycemia is known to activate NADPH oxidases, impair antioxidant enzyme activity, and drive mitochondrial dysfunction^6,15^.

### Metabolic dysfunction and energy depletion in wounds of diabetic minipig

Diabetic wounds also showed downregulation of genes involved in various metabolic pathways, including branched-chain amino acid (**BCAA**) catabolism (IVD, 1CD2097A1, OXCT1, MMUT, and ACADSB). This reduction leads to BCAA accumulation and increased production of their ketoacid byproducts (**BCAK**s), thereby exacerbating insulin resistance, mitochondrial dysfunction, and ROS generation, while also impairing ATP generation, which is essential for cell proliferation, migration, and collagen synthesis^16,17^.

Similarly, downregulation of key genes associated with fatty acid (**FA**) metabolism (ELOVL5, THRSP, ACLY, ACSL1, SCD, ELOVL6, PON3, ACACA, CYP1A1, CYP4B1, MORC2, CYP2U1, FADS1, ACSL3, MMUT) and peroxisome proliferator-activated receptor (**PPAR**) signaling (OLR1, ME1, ACSL1, SCD, SLC27A6, FABP3, LPL, and ACSL3) indicates impaired lipogenic capacity and energy substrate utilization in diabetic wounds. Given that FA oxidation (**FAO**) is critical for efficient ATP production, reduced expression of FAO-related genes likely compromises the energetic capacity of cells in diabetic wounds^17^. Importantly, FAO is critical for efficient oxidative phosphorylation and is therefore a key factor in the phenotypic transition of macrophages from pro-inflammatory M1, a phenotype reliant on glycolysis, to pro-resolution M2, which requires oxidative phosphorylation^12,17^. Chronic downregulation of FAO-related genes may prevent M1-to-M2 macrophage polarization. Indeed, CCR7 immunostaining identified increased pro-inflammatory (CCR7+) macrophage accumulation, while pro-resolution (CD206+) macrophages were substantially decreased in diabetic wounds compared to non-diabetic wounds (**Figure 2E;Figure S2B-C**), consistent with impaired M1 to M2 macrophage polarization known to occur in hyperglycemic environments and characteristic of human DFUs. Importantly, M1 macrophages perpetuate pro-inflammatory cytokine production (e.g., TNF) and ROS generation, thereby sustaining inflammation and oxidative stress^12,17^.

These gene expression changes are consistent with essential roles of BCAA and FA metabolism in wound healing and inflammation, identifying potential molecular targets for further exploration.

### ECM organization dysfunction in wounds of diabetic minipigs

Genes associated with ECM production and organization were robustly downregulated, including collagen isoforms (COL8A1, COL9A2, COL15A1, COL25A1), crosslinking enzymes (COLGALT2, P4HA1, PLOD2), matricellular proteins (TGFB3, FBN1, SPP1), and mediators of cell-ECM interactions (LAMA1, LAMA3, ACAN, ITGA8). Downregulation of these components may result in an insufficient, immature, poorly cross-linked matrix, adversely affecting cell migration and tissue integrity. Diminished and disorganized ECM has been previously noted in human DFUs and is thought to affect cell-matrix signaling required for fibroblast and keratinocyte migration and differentiation^5,17,18^. These transcriptomic results align with the histological observations of reduced collagen deposition and altered tissue architecture in the diabetic wounds (**Figure 1F**).

## Conclusion

In the present work, we transcriptionally profiled a preclinical diabetic wound-healing model in Yucatan minipigs. Induction of diabetes by STZ resulted in diabetic wounds that exhibited delayed rates of wound closure compared to non-diabetic wounds. Transcriptomic analysis identified enriched autoimmune pathways, elevated oxidative stress, dysregulated metabolic pathways, and comprehensive suppression of ECM organization. These findings were supported by histological findings of increased M1 macrophage polarization, increased leukocyte content, and decreased ECM deposition in diabetic wounds. Our findings confirm that many of the pathophysiological and molecular hallmarks of human DFUs are faithfully recapitulated in this diabetic porcine model. Ultimately, anatomical similarity between pig and human skin, and robust generation of diabetic wound pathology position this model as a valuable preclinical platform for testing emerging therapeutics in a physiologically relevant context. Together, the gene signatures identified here point to tractable pathways that may contribute to pathological wound healing in diabetic patients, providing a resource to be further mined for new therapeutic targets.

## Supporting information

Supplementary Material, Methods, and Results

## Acknowledgments

This work was supported by the National Institutes of Health (NIH) (R01 EB02869, R01 EB019409, and T32 DK101003) and conducted in collaboration with Pluris Research Inc. (Franklin, TN, USA). We also acknowledge the Translational Pathology Shared Resource supported by NCI/NIH Cancer Center Support Grant P30CA068485.

## Data Availability Statement

All data generated in this study are available within the article or Supporting Information or from the corresponding author upon reasonable request. Gene expression data generated and analyzed in this study are available in the Gene Expression Omnibus (GEO) repository, accession number GSE315429.

