## Supplementary Material, Methods, and Results for "Transcriptomic Profiling of Diabetic Porcine Wound Healing Model Identifies Key Metabolic, Inflammatory, and Oxidative Stress Pathways"

**Supplementary Information**

*Materials and Methods*

Induction and maintenance of diabetes – All procedures were reviewed and approved by Pluris Research’s Institutional Animal Care and Use Committee (IACUC) under protocol VU1. Methods for the induction of diabetes using streptozotocin were adapted from previous reports^1-4^. Briefly, a female Yucatan minipig (20-24 kg, 6 months of age) was anesthetized and maintained under 2-3% isoflurane. A sterile streptozotocin solution (100 mg/mL in sodium citrate buffer) was infused intravenously at a dose of 125 mg/kg over 15 minutes. Following induction, a continuous blood glucose monitor (CGM) (Dexcom G7, Dexcom, Inc.) was placed on the posterior cervical region of the animal. The CGM was replaced every ten days, or earlier if necessary, prior to initiation of the wound healing study and was continued throughout the entire duration of the wound healing study. The minipig was provided with a 10% sucrose solution in water for the initial 48 hours following induction to prevent hypoglycemia upon beta cell death and insulin release. Subcutaneous insulin (Lantus, Sanofi Group) was administered daily (0.3 u/kg), beginning one week after induction. The minipig was maintained for sixty days before initiating the wound-healing study, with insulin administration ending one week before wounding.

Porcine excisional wound healing model – Both diabetic and non-diabetic minipigs were subject to identical procedures for creating full-thickness excisional wounds. Briefly, female Yucatan minipigs were anesthetized and maintained under 2-3% isoflurane for the duration of surgery. The dorsal skin was shaved and disinfected with 70% ethanol and Betadine. Twenty-eight full-thickness wounds were created on the dorsum of the minipigs using a 2 cm biopsy punch (Radha Meditech), each spaced 2 cm apart. Wounds were oriented in two parallel rows, each containing seven wounds, one on either side of the spine. Each wound was covered with a Tegaderm dressing (3M) and surrounded by 3M Reston™ foam padding (3M), before being covered with a Tendersorb™ underpad (Covidien) and MediChoice tubular net gauze (Owens & Minor). Following surgery, dressings were changed every 2 to 3 days (POD 1, 3, 6, 8, and 10), under anesthesia. Just prior to euthanasia, wound samples were collected from anesthetized pigs using 2 cm biopsy punches for bulk RNA sequencing analysis. The minipigs were then euthanized via intracardiac administration of Euthasol (Virbac), and the remaining wound tissue was excised for histological assessment.

Wound area quantification – Digital photographic images of each wound, with a calibrated scale, were acquired during surgery and all subsequent dressing changes. The wound was quantified in ImageJ and normalized with respect to the initial wound area on the operative day. Statistical significance was determined at each time point using a Welch’s t test in GraphPad Prism 10.

RNA sequencing and data analysis – Excised tissue from biopsy punches was stored in RNAlater™ Stabilization Solution (Invitrogen) at -80 °C until the time of RNA isolation. Total RNA was extracted by lysing tissue in TRIzol™ Reagent (Invitrogen) using a TissueLyser II (QIAGEN). The tissue homogenate was phase-separated with chloroform, and the aqueous RNA phase was further purified using a RNeasy Mini Kit (QIAGEN) per the manufacturer’s recommendations. Total RNA was sent to Novogene for quantification, quality control, library preparation, sequencing, and analysis. Briefly Bulk RNA-seq was conducted using the Illumina NovaSeq X Plus Series platform. Raw reads were processed with fastp software to remove adaptor sequences and retain clean reads. Clean reads were mapped against the Sus scrofa reference genome (Sscrofa11.1) using HISAT2 (version 2.2.1). Mapped reads were then assembled using StringTie (version 2.2.3) and quantified using featureCounts (version 2.0.6). Differential gene expression analysis was performed on raw counts using DESeq2 (version 1.42.0) using the Benjamini-Hochberg method to calculate the false discovery rate (FDR) and adjusted p-value. Differentially expressed genes (DEGs) were defined as having a |Log2FC| ≥ 0.5 and adjusted p-value < 0.05. Functional enrichment analysis was conducted using clusterProfiler^5^ (version 4.10.1), with gene sets imported from the MSigDB collections using the geneset R package (version 0.2.7). All enrichment analysis was corrected using the Benjamini-Hochberg method to calculate the false discovery rate (FDR) and adjusted p-value. Heatmaps, volcano plots, and KEGG pathway diagrams were made using ComplexHeatmap (version 2.18.0), genekitr (version 1.2.8), and pathview (version 1.42.0), respectively.

Histology and immunohistochemistry – Excised tissue was fixed in 10% neutral buffered formalin, processed, and embedded in paraffin. Histology was carried out as previously described^6^. Briefly, tissue sections (5 µm) were deparaffinized and rehydrated through gradients of xylene, histological grade alcohols, and tris-buffered saline/0.1% Tween-20 (TBST). Gomori’s trichrome and hematoxylin and eosin (H&E) were performed per the manufacturer's recommendations. Antigen retrieval was conducted in LabVision™ citrate heat-induced epitope retrieval solution (Epredia) for 20 minutes at 96 °C in a pretreatment (PT) module (Epredia). The sections were incubated with 3% hydrogen peroxide in TBST for 40 minutes and then incubated with protein block (Dako) for 20 minutes. The sections were then incubated with their respective primary antibodies (mouse anti-8-hydroxy-2’-deoxyguanosine (ab48508 Abcam, 1:2000), rabbit anti-CCR7 (ab32527 Abcam, 1:1000), rabbit anti-CD206 (ab91279 Abcam, 1:1000), rabbit anti-myeloperoxidase (A039829-2 Dako, 1:1000), or mouse anti-cytokeratin14 (MCA890 Bio-Rad, 1:2000)) for 60 minutes at room temperature. The sections were then immediately incubated with the corresponding secondary antibodies (donkey anti-mouse HRP (Envision+/HRP, Dako) for mouse primary antibodies or donkey anti-rabbit HRP (Envision+/HRP, Dako) for rabbit primary antibodies) for 30 minutes at room temperature and incubated with 3,3′-diaminobenzidine (DAB) substrate (Dako) for 10 minutes. Slides were then rinsed with TBST buffer, counterstained in hematoxylin for 2 minutes, rinsed with running DI water, dehydrated, and mounted with Acrytol mounting media.

Immunohistological (IHC) sections were analyzed using QuPath software. Briefly, slides for 8-OHdG, CCR7, and CD206 were color deconvolved, and the number of brown nuclei (positive for immunoreactivity with DAB) was normalized to the total number of nuclei within a field of view. A minimum of three fields of view per sample were analyzed to obtain an average quantification for each sample. Percent re-epithelialization was calculated as the percentage of the wound covered by an epidermal layer with nuclei positive for cytokeratin 14 (CK14).

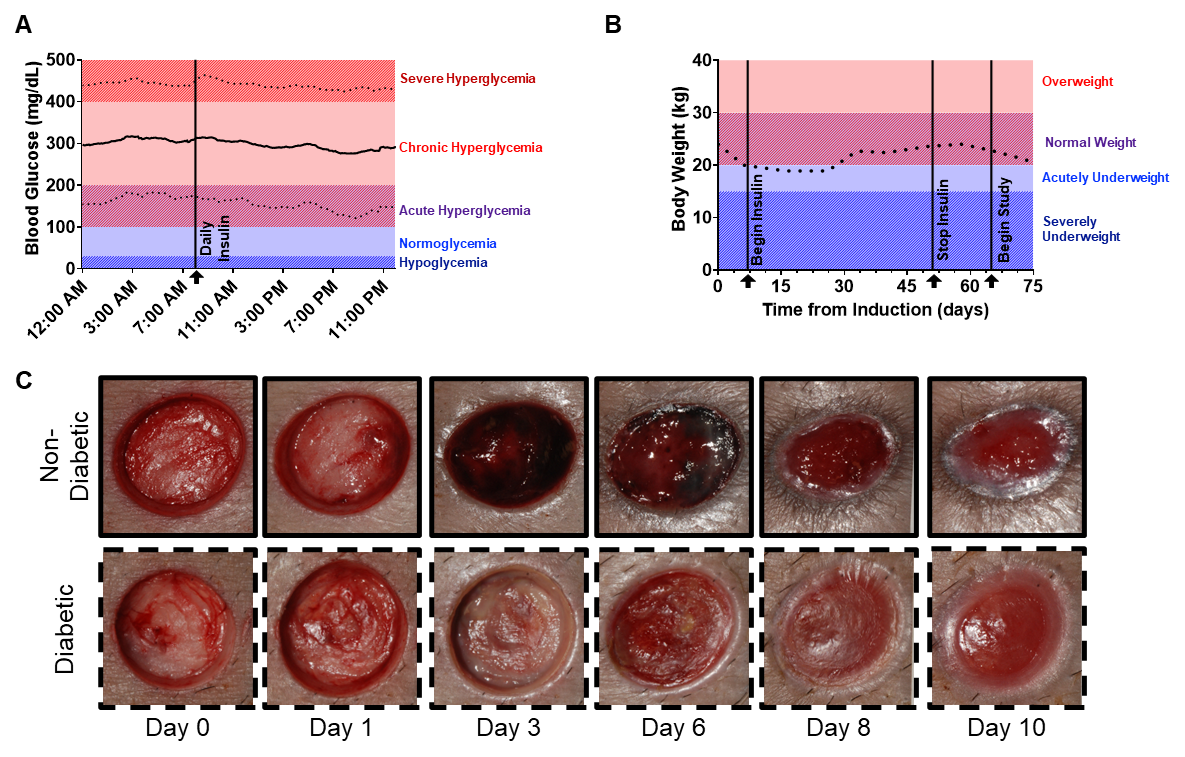
Figure S1. Streptozotocin induction and wound healing study in Yucatan minipig. **(A)** Average (mean ± std.) temporal daily blood glucose (mg/dL) and **(B)** body weight of diabetic minipig over the course of the study. **(C)** Representative images of wounds at day 0, 1, 3, 6, 8, and 10.

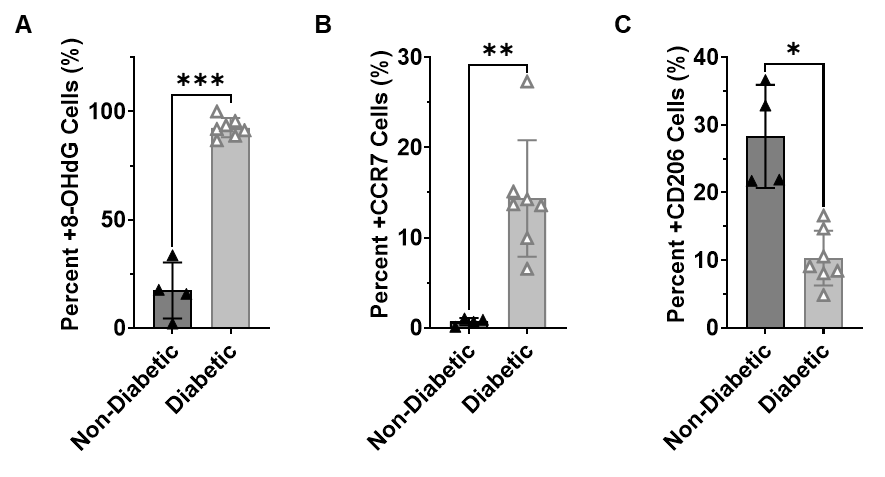

Figure S2. Quantification of **(A)** 8-OHdG, **(B)** CCR7, and **(C)** CD206 immunohistochemistry as a percentage of positive cells in the wound bed. *P<0.05, **P<0.01, and ***P<0.001, by Welch’s unpaired t-test.

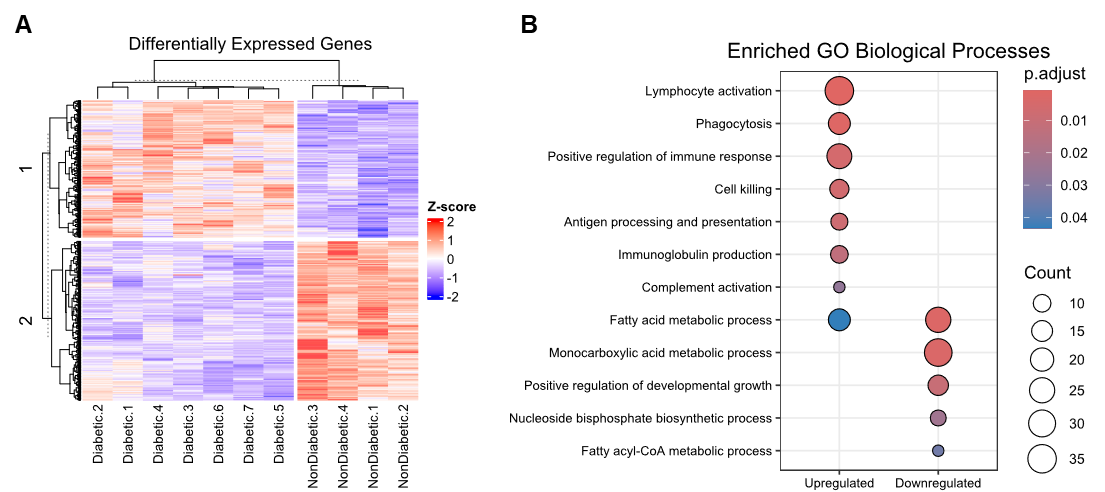

Figure S3. Differential gene expression and functional enrichment analysis. **(A)** Clustered heatmap of differentially expressed genes (DEGs), with row cluster 1 corresponding to upregulated genes in diabetic wounds and row cluster 2 corresponding to downregulated genes. **(B)** Gene ontology biological processes enriched within diabetic wounds.

Figure S4. KEGG enrichment – Graft versus host disease. **(A)** Pathway schematic illustrating genes that were upregulated (red) within the pathway in diabetic vs non-diabetic wounds. **(B)** Clustered heatmap of upregulated genes within the graft versus host disease pathway.

**B**

**A**

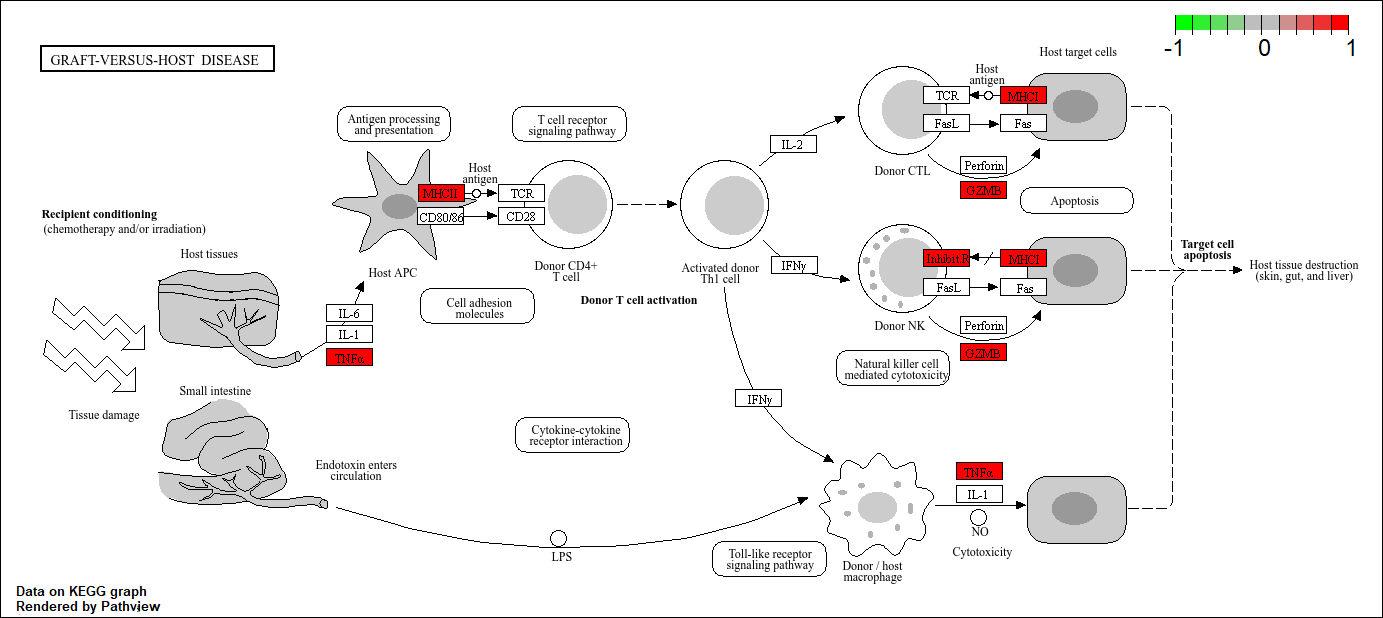

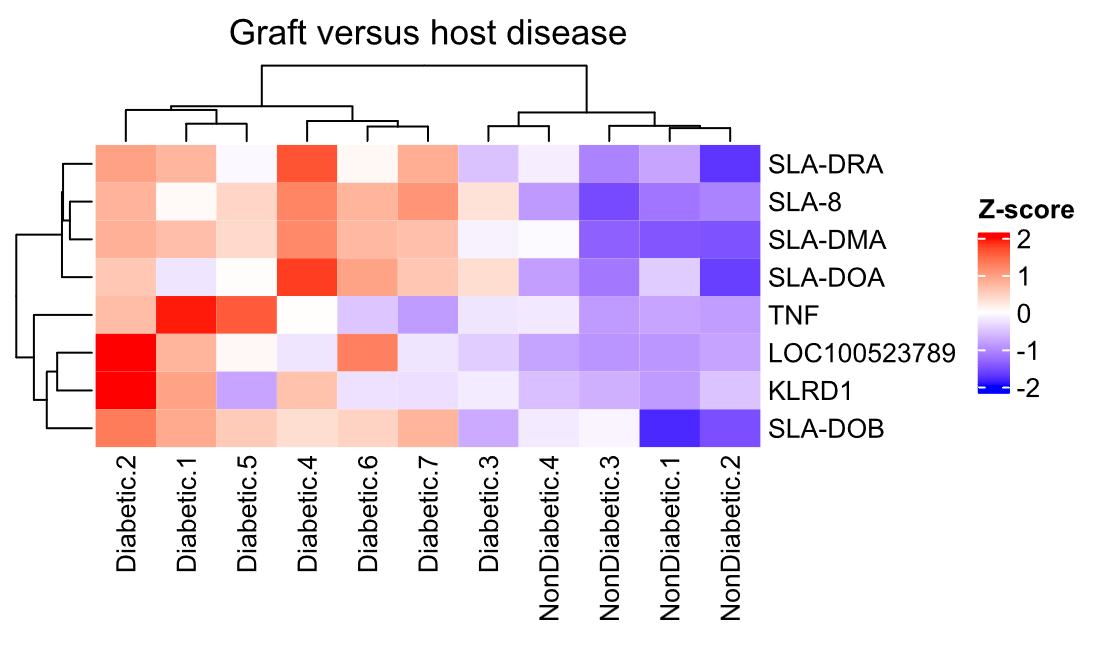

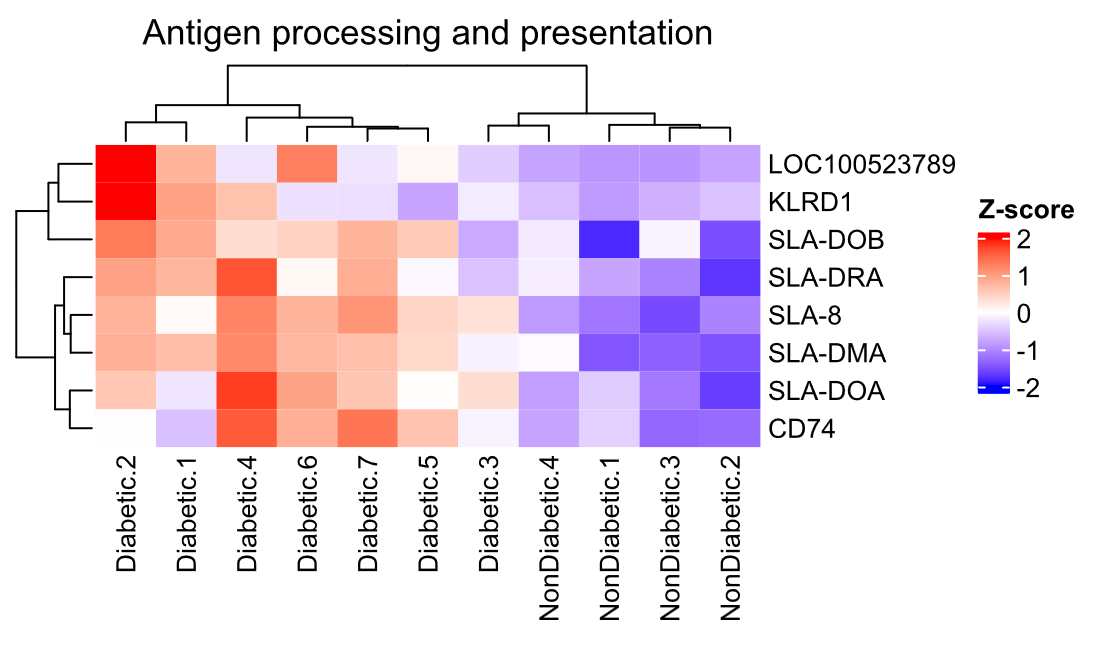

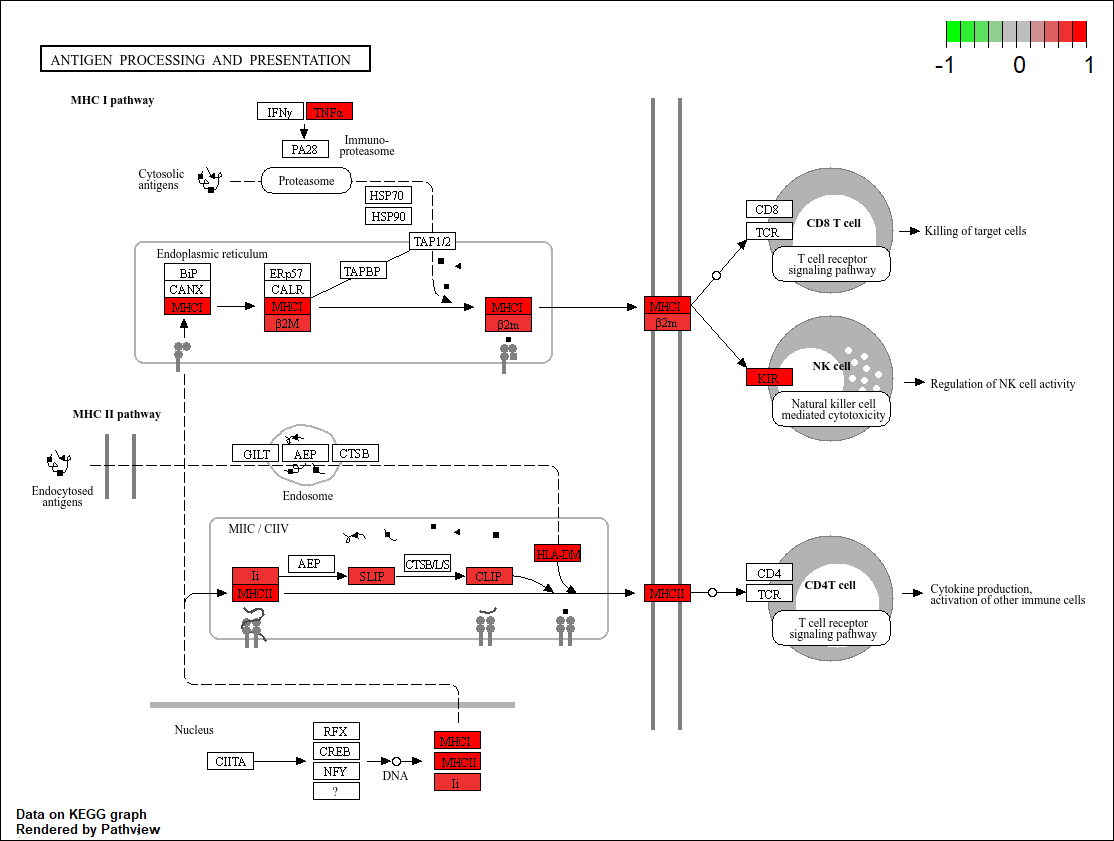
Figure S5. KEGG enrichment – Antigen processing and presentation. **(A)** Pathway schematic illustrating genes that were upregulated (red) within the pathway in diabetic vs non-diabetic wounds. **(B)** Clustered heatmap of upregulated genes within the antigen processing and presentation pathway.

**B**

**A**

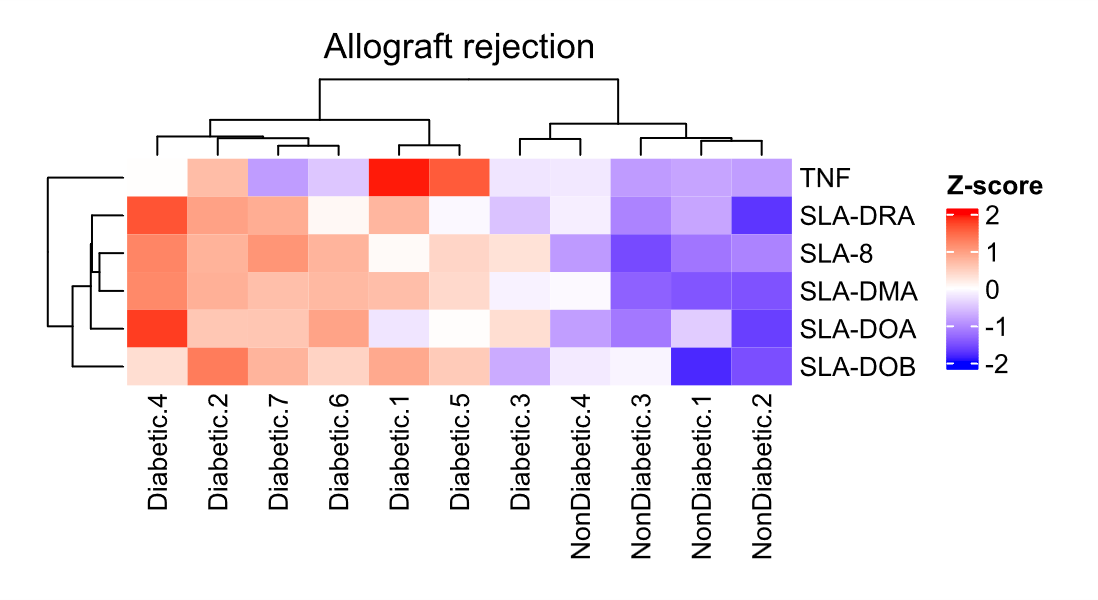
Figure S6. KEGG enrichment – Allograft rejection. **(A)** Pathway schematic illustrating genes that were upregulated (red) within the pathway in diabetic vs non-diabetic wounds. **(B)** Clustered heatmap of upregulated genes within the allograft rejection pathway.
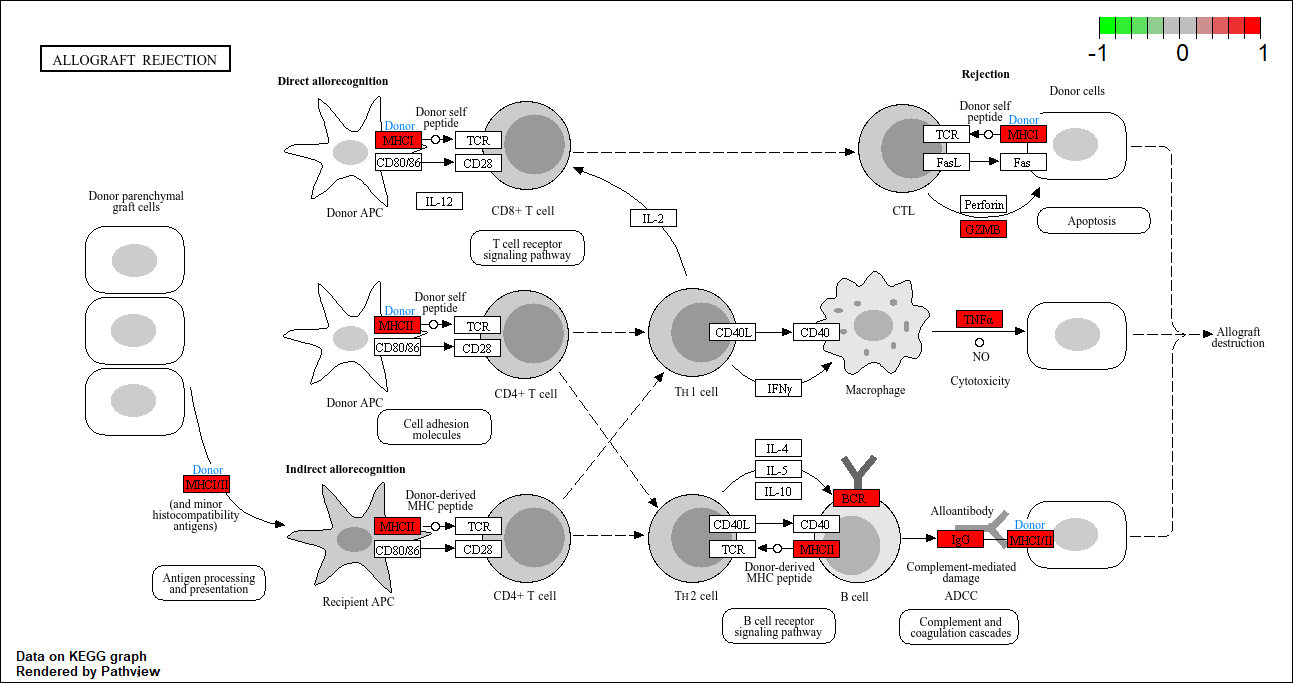

**B**

**A**

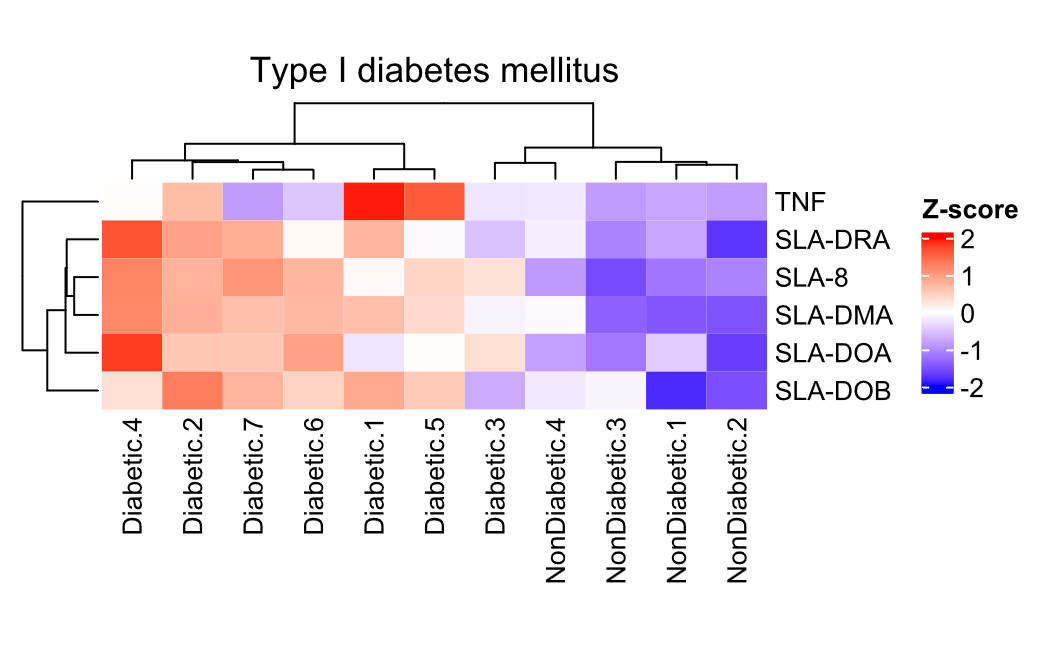

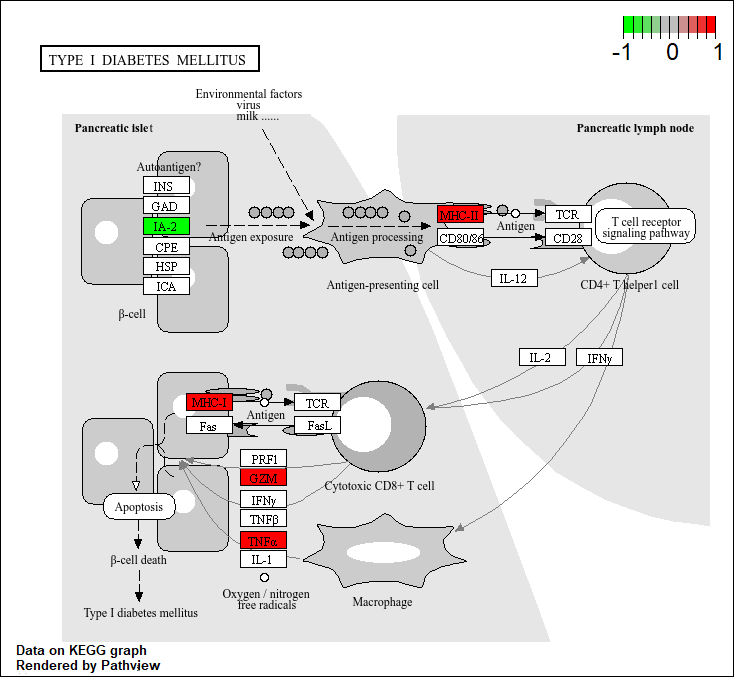
Figure S7. KEGG enrichment – Type I diabetes mellitus. **(A)** Pathway schematic illustrating genes that were upregulated (red) and downregulated (green) within the pathway in diabetic vs nondiabetic wounds. **(B)** Clustered heatmap of upregulated genes within the type I diabetes mellitus pathway.

**A**

**B**

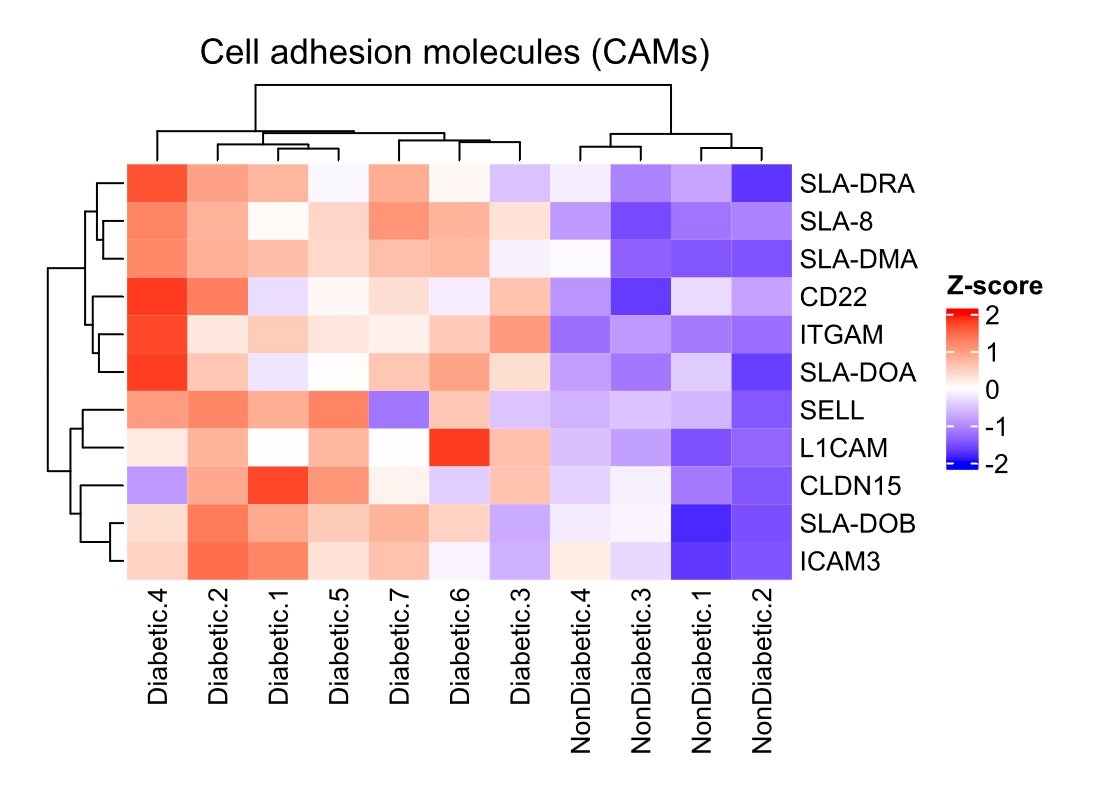
Figure S8. KEGG enrichment – Cell adhesion molecules (CAM) interactions. **(A)** Pathway schematic illustrating genes that were upregulated (red) and downregulated (green) within the pathway in diabetic vs nondiabetic wounds. **(B)** Clustered heatmap of upregulated genes within the cell adhesion molecules (CAM) interactions pathway.
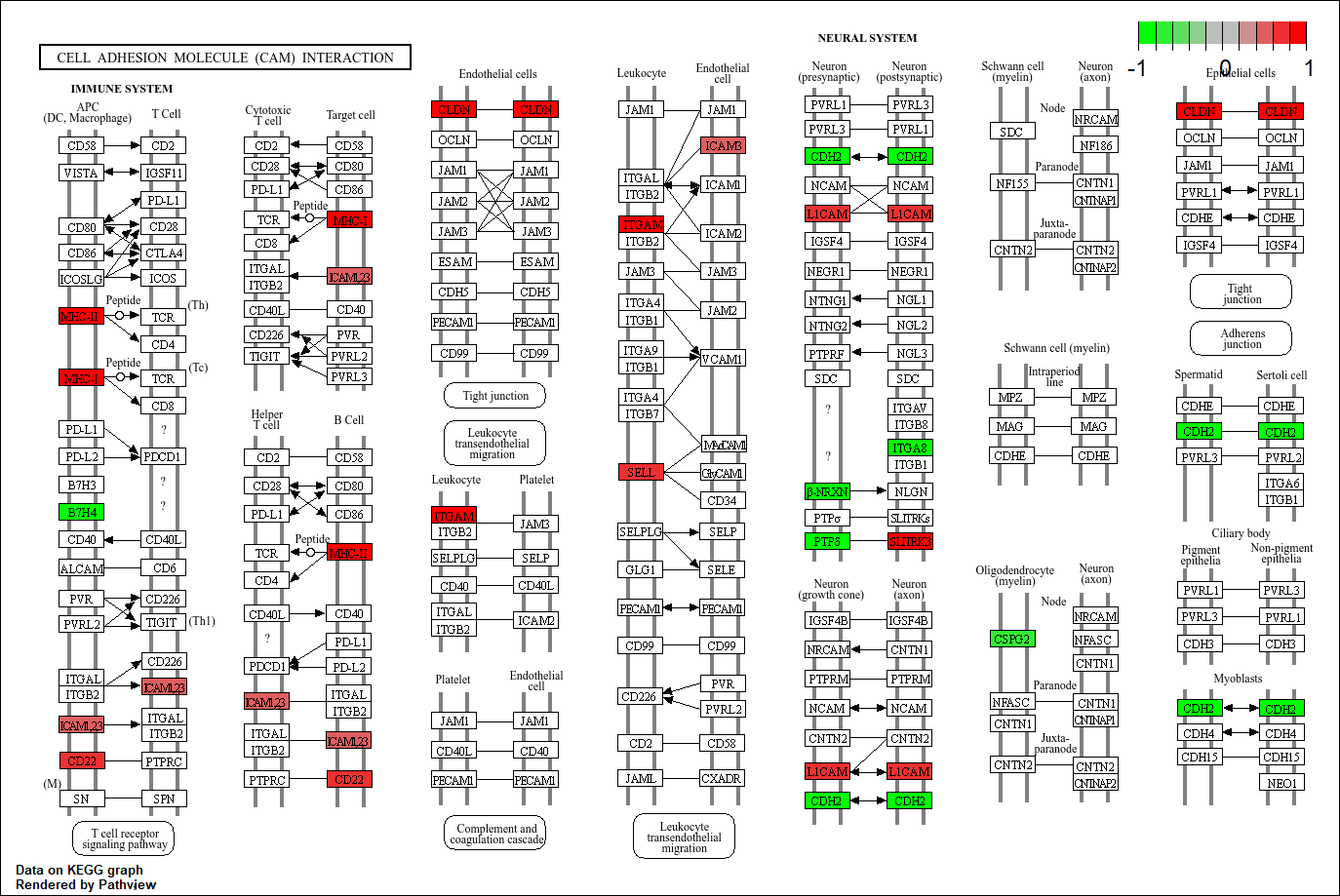

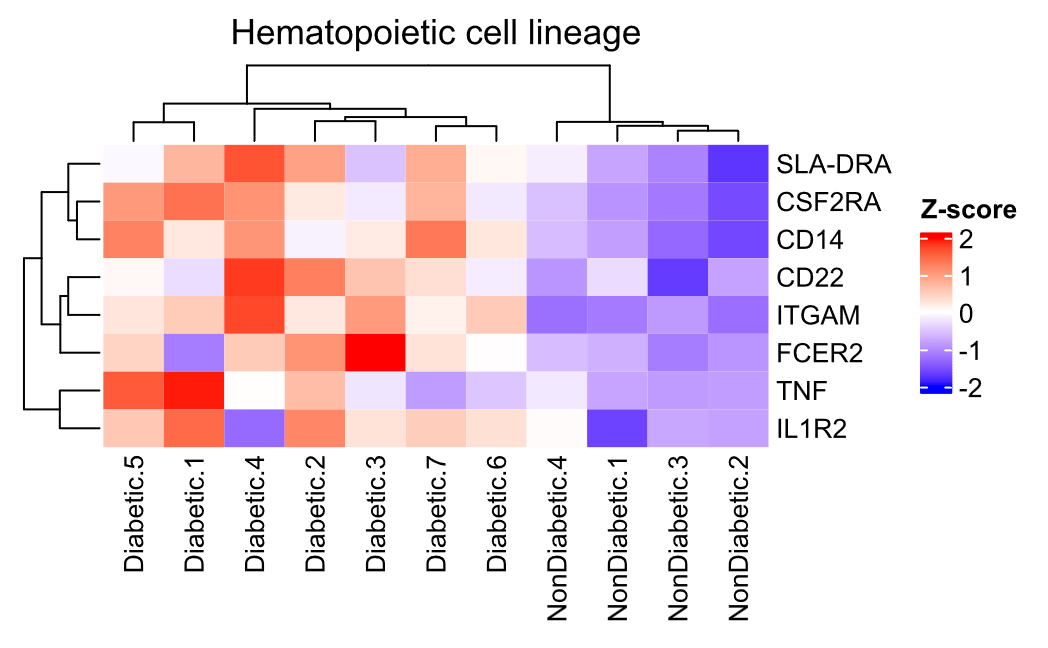
Figure S9. KEGG enrichment – Hematopoietic cell lineage. **(A)** Pathway schematic illustrating genes that were upregulated (red) and downregulated (green) within the pathway in diabetic vs nondiabetic wounds. **(B)** Clustered heatmap of upregulated genes within the hematopoietic cell lineage pathway.
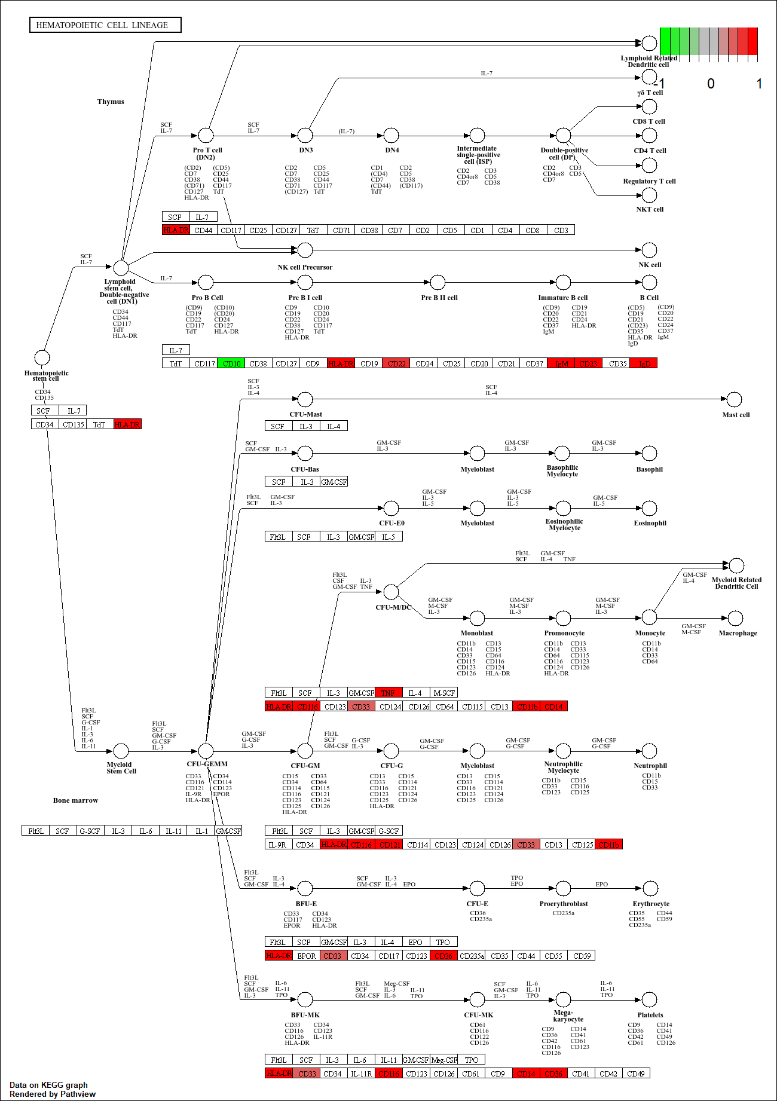

**B**

**A**

**B**

**A**

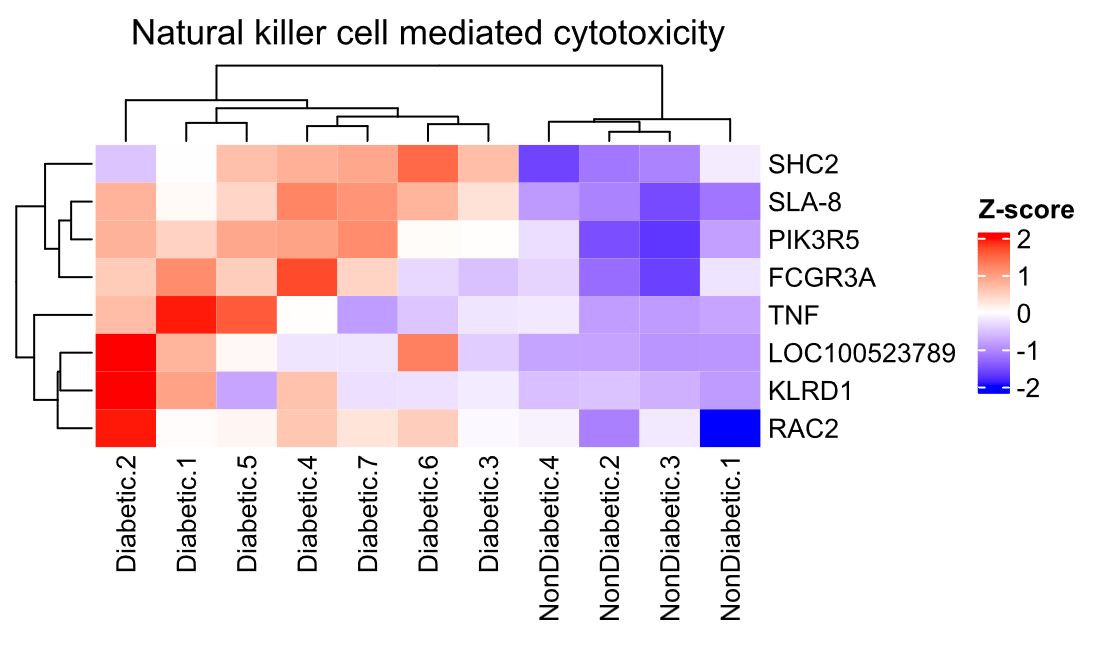
Figure S10. KEGG enrichment – Natural killer cell mediated cytotoxicity. **(A)** Pathway schematic illustrating genes that were upregulated (red) within the pathway in diabetic vs nondiabetic wounds. **(B)** Clustered heatmap of upregulated genes within the natural killer cell mediated cytotoxicity pathway.
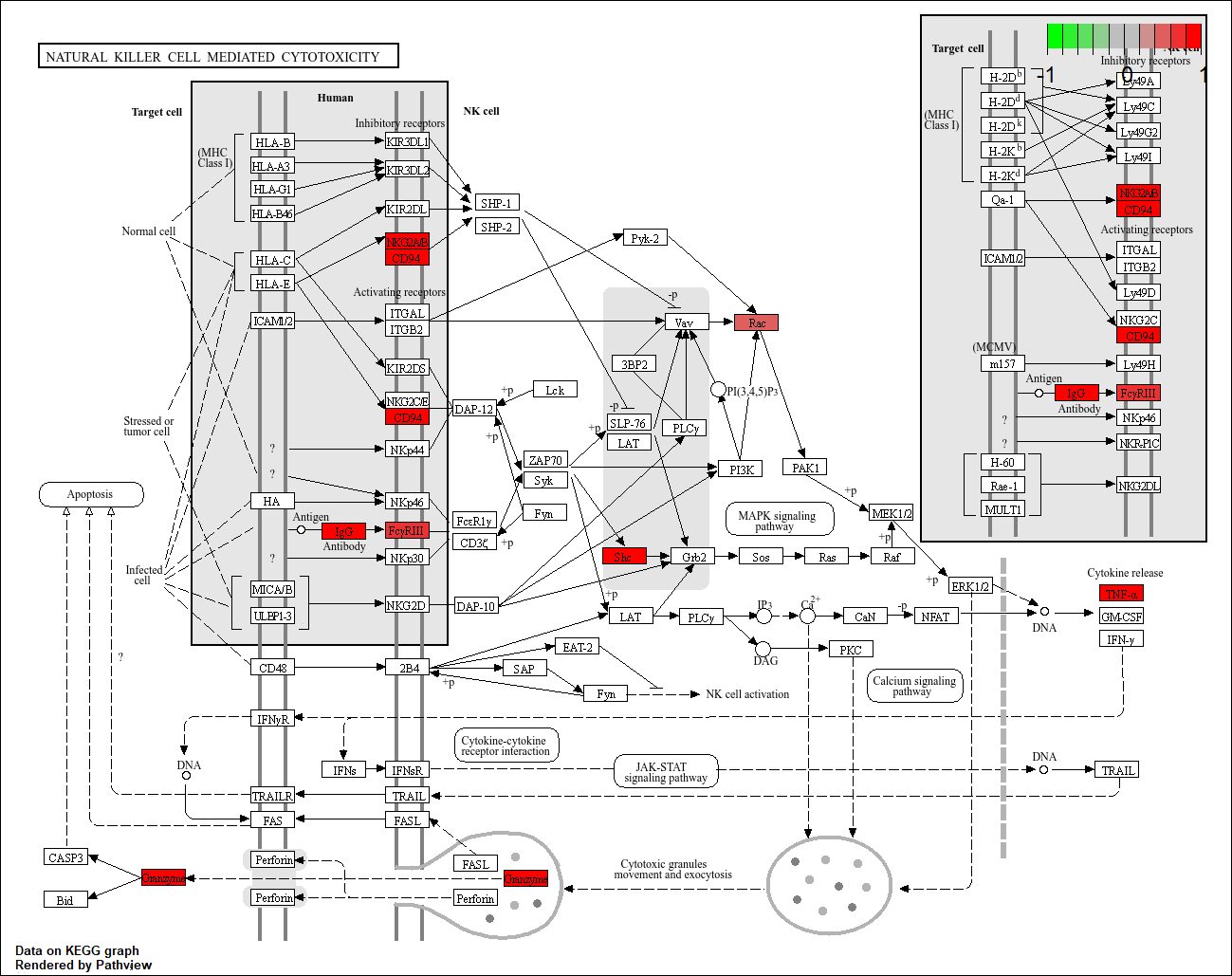

**B**

**A**

Figure S11. KEGG enrichment – Arachidonic acid metabolism. **(A)** Pathway schematic illustrating genes that were upregulated (red) and downregulated (green) within the pathway in diabetic vs nondiabetic wounds. **(B)** Clustered heatmap of upregulated genes within the arachidonic acid metabolism pathway.
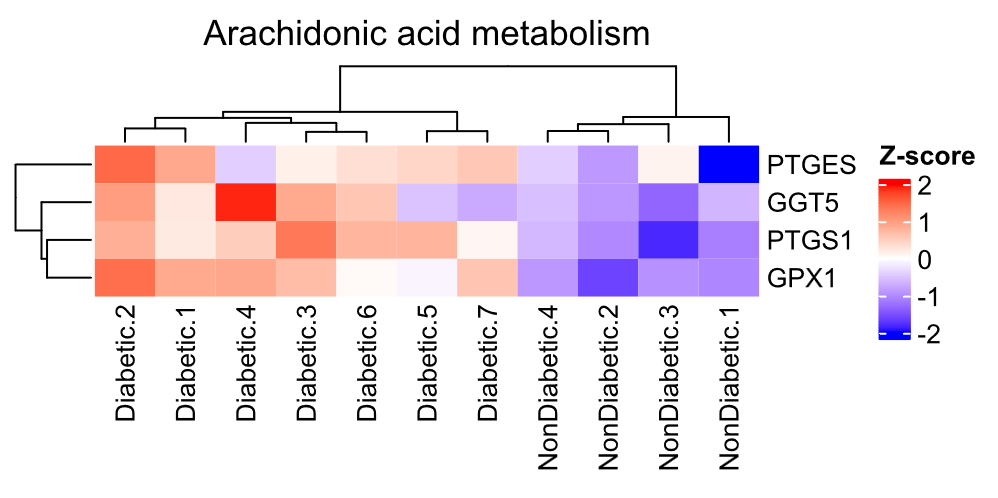

**B**

**A**

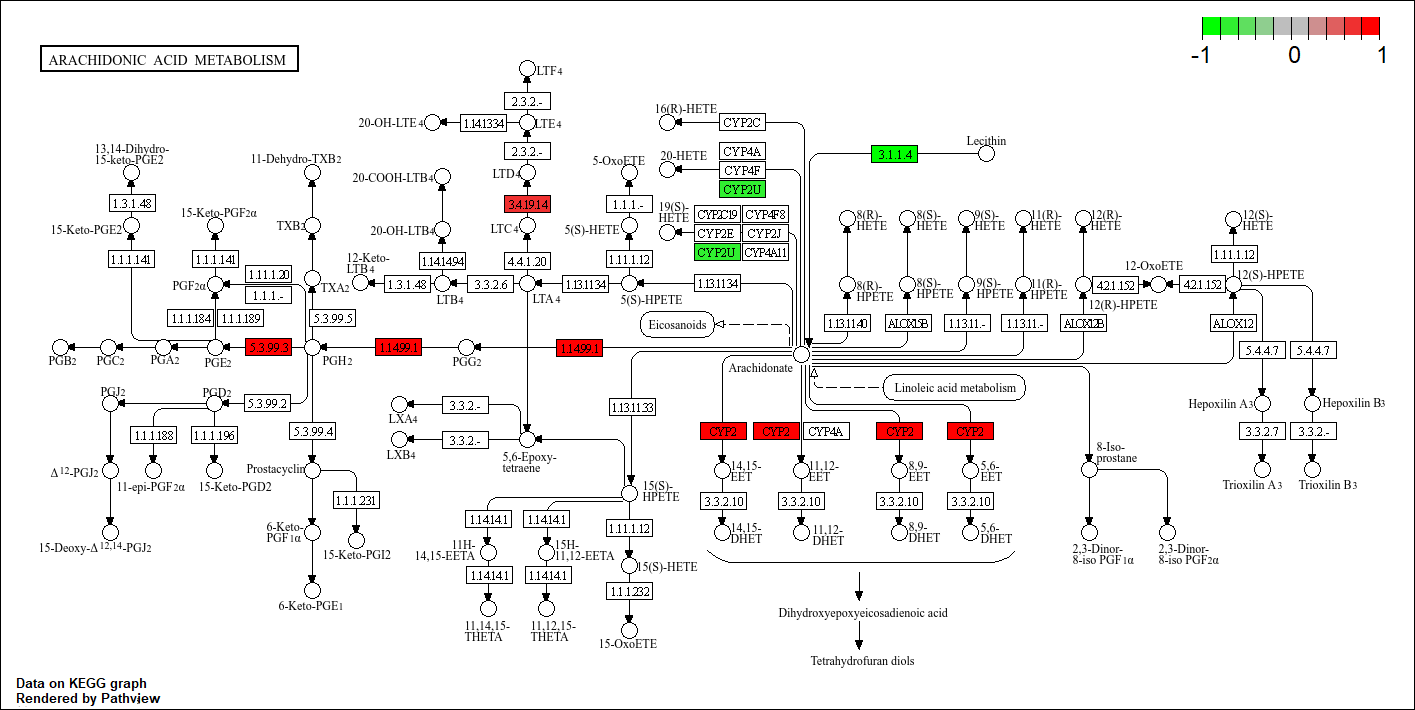

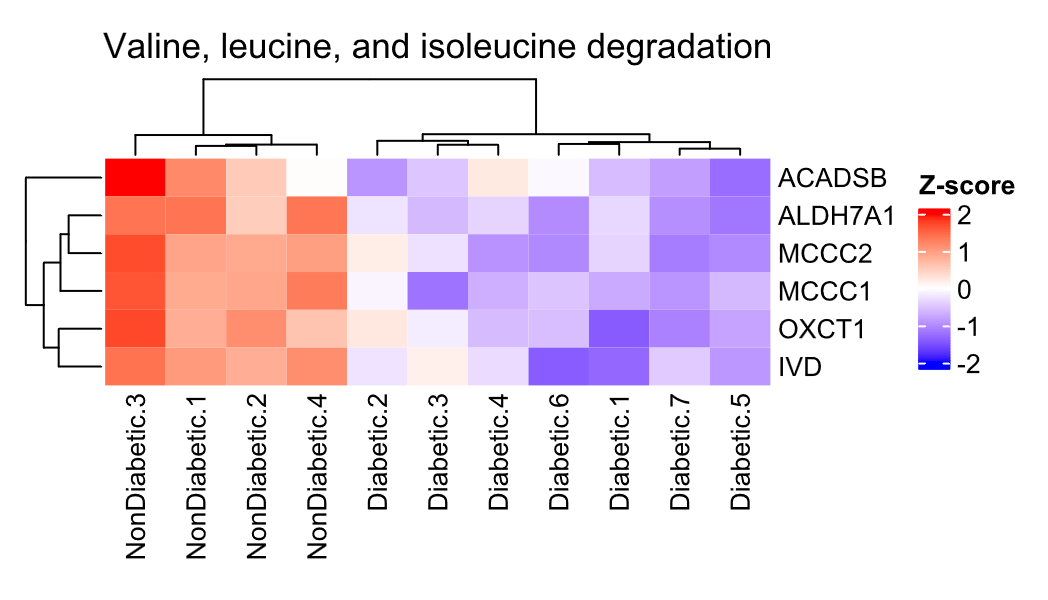
Figure S12. KEGG enrichment – Valine, leucine, and isoleucine degradation. **(A)** Pathway schematic illustrating genes that were downregulated (green) within the pathway in diabetic vs nondiabetic wounds. **(B)** Clustered heatmap of downregulated genes within the Valine, leucine, and isoleucine degradation pathway.
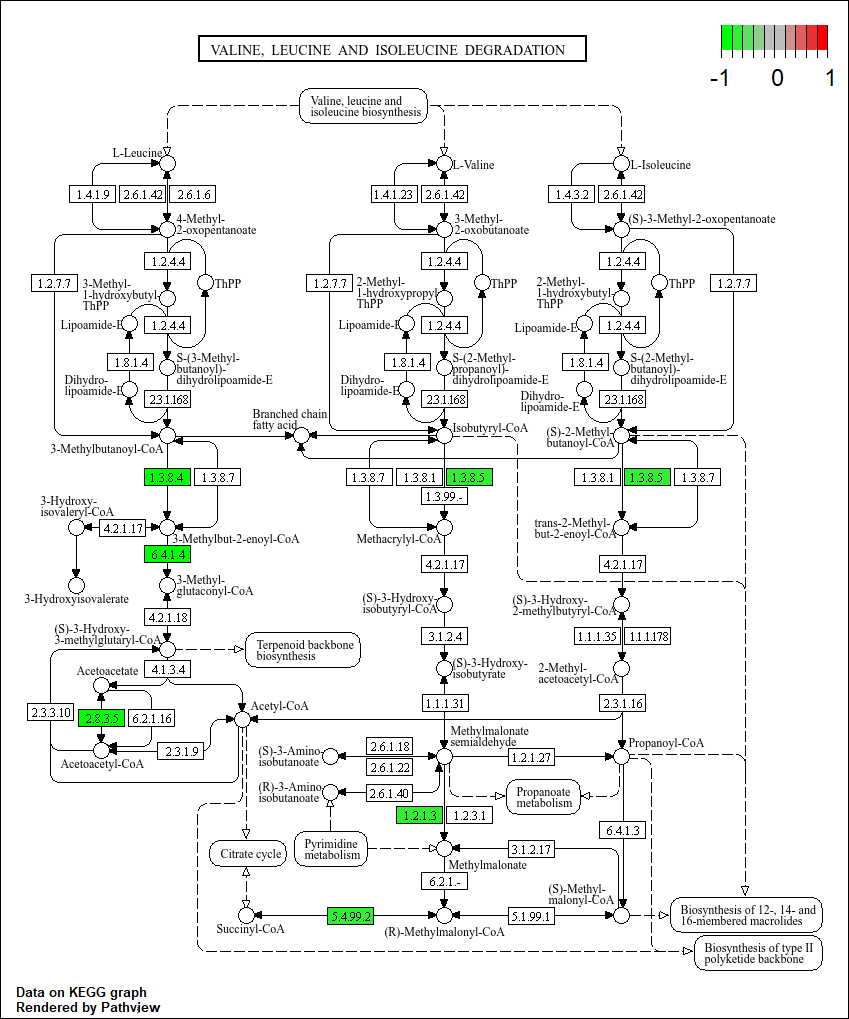

**B**

**A**

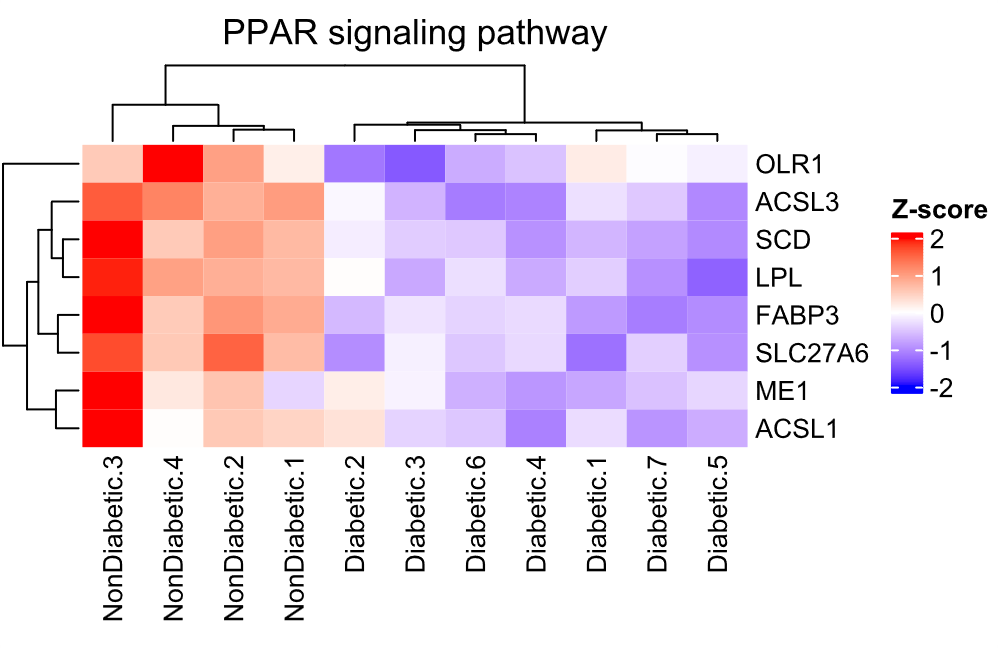
Figure S13. KEGG enrichment – PPAR signaling pathway. **(A)** Pathway schematic illustrating genes that were upregulated (red) and downregulated (green) within the pathway in diabetic vs nondiabetic wounds. **(B)** Clustered heatmap of downregulated genes within the PPAR signaling pathway.
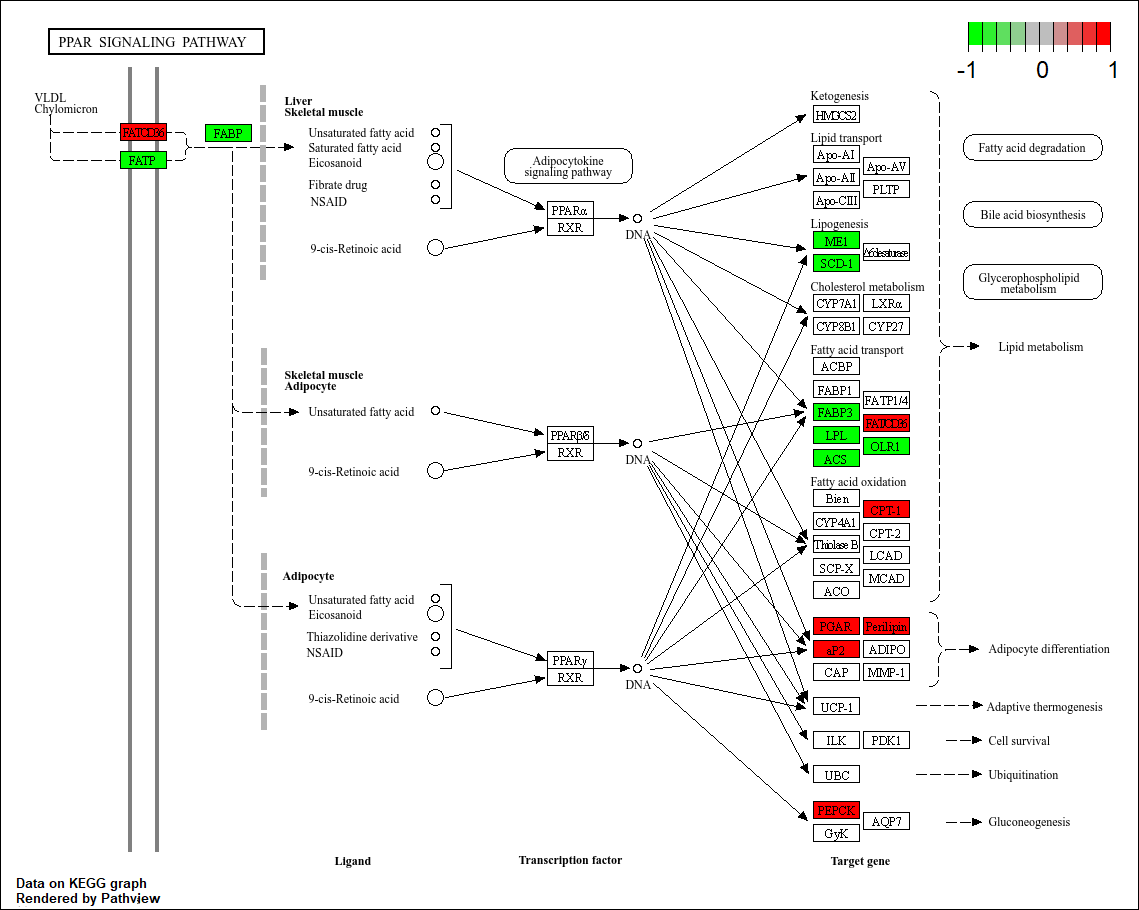

**B**

**A**

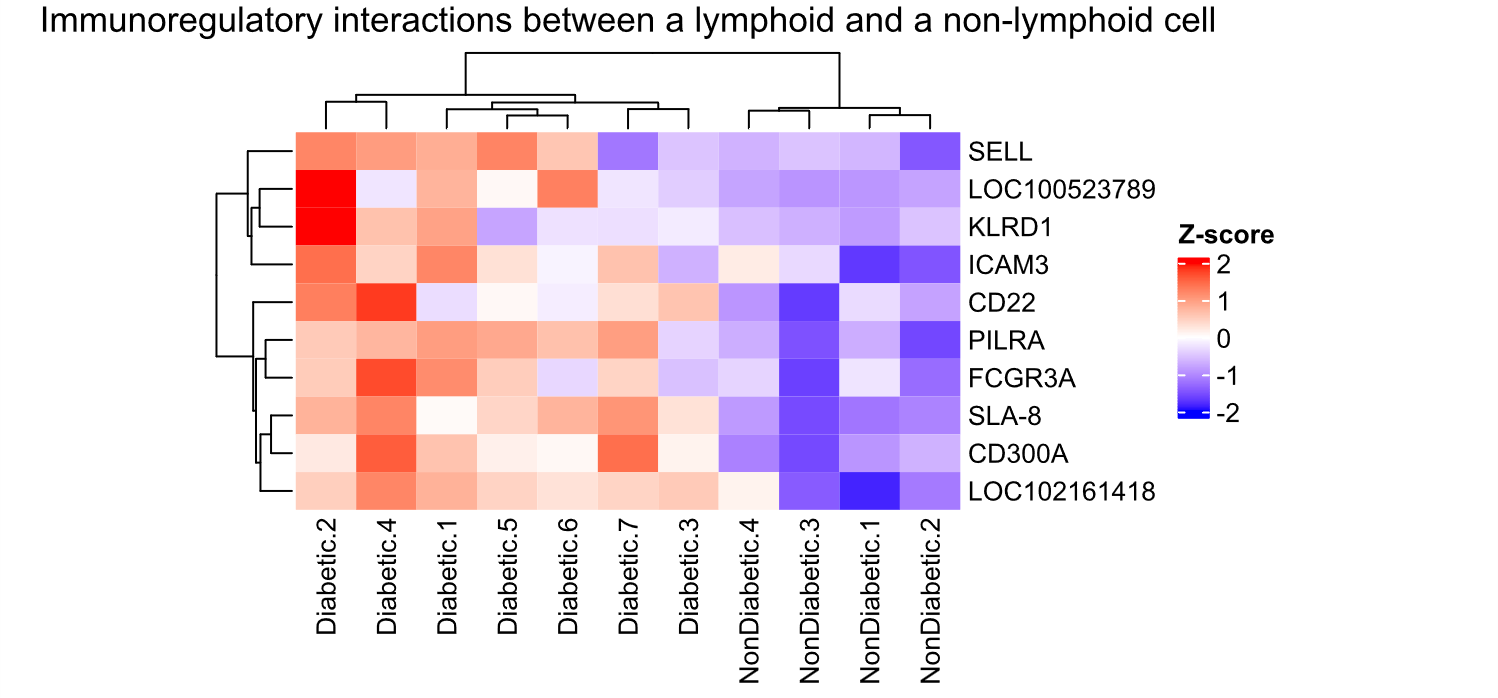

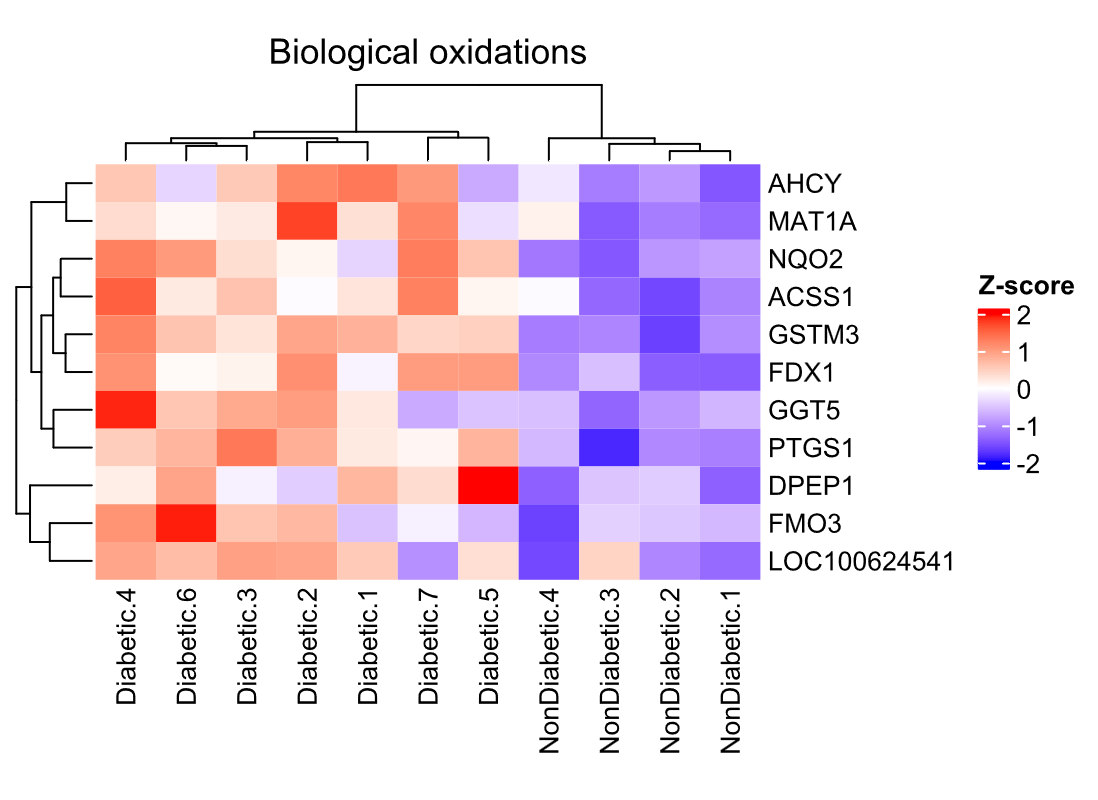
Figure S14. Upregulated Reactome enrichment. **(A)** Clustered heatmap of genes upregulated within the immunoregulatory interactions between lymphoid and non-lymphoid cell pathway in diabetic relative to nondiabetic wounds. **(B)** Clustered heatmap of genes upregulated within the biological oxidations pathway in diabetic relative to nondiabetic wounds.

**A**

**B**

Figure S15. Downregulated Reactome enrichment – Clustered heatmap of genes downregulated in diabetic relative to nondiabetic wounds within the **(A)** fatty acyl-CoA biosynthesis, **(B)** branched chain amino acid catabolism, **(C)** fatty acid metabolism, and **(D)** defects in vitamin and cofactor metabolism pathways.

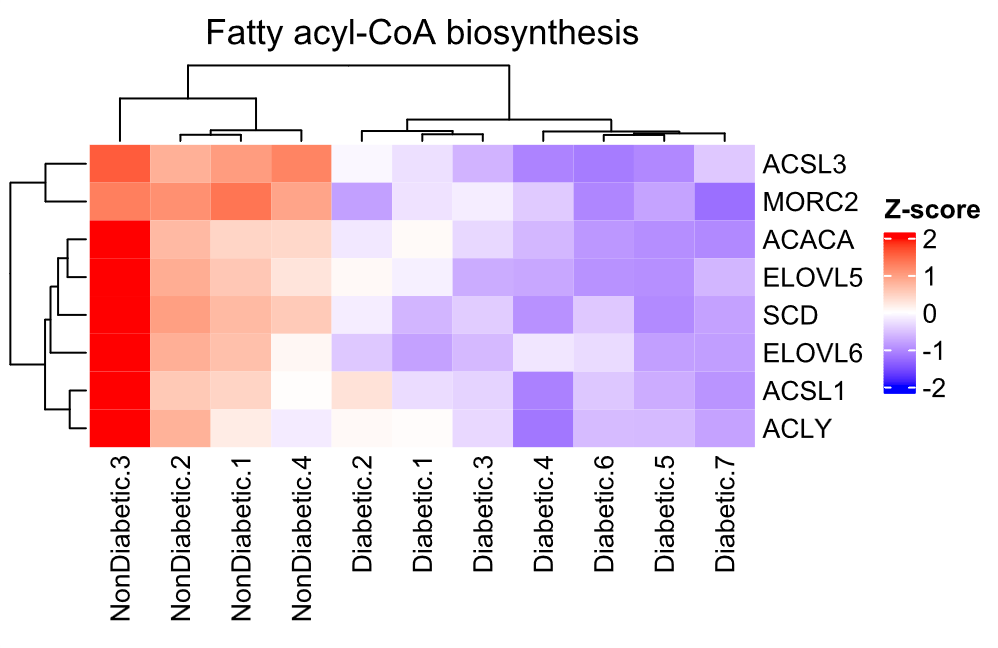

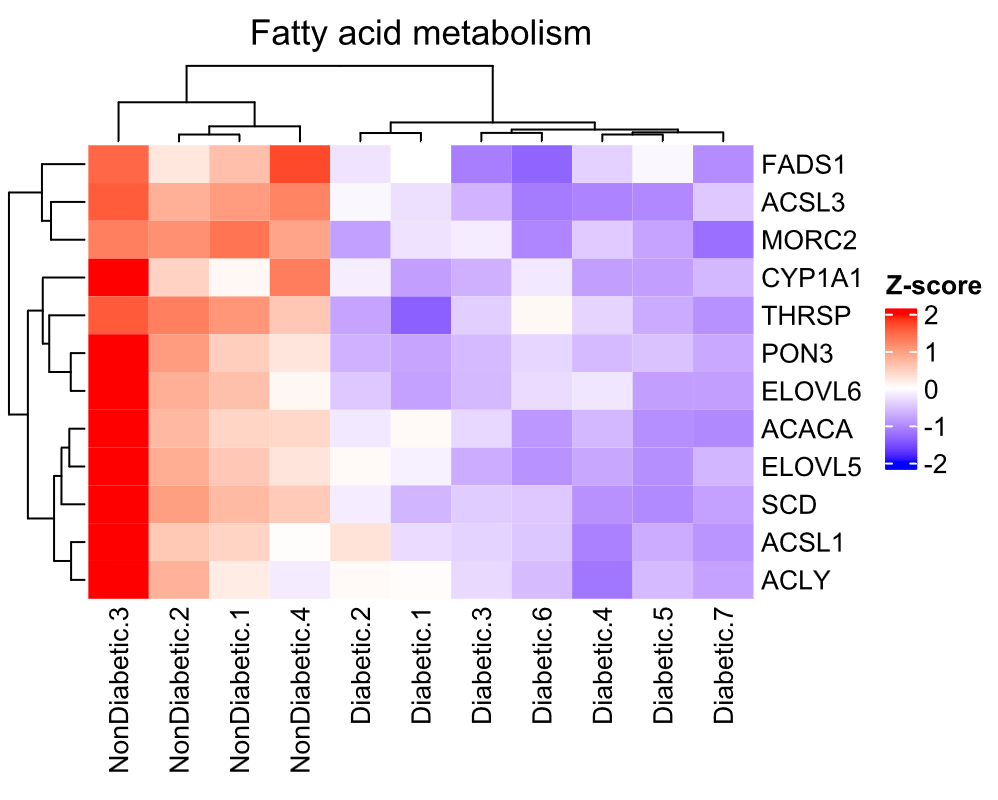

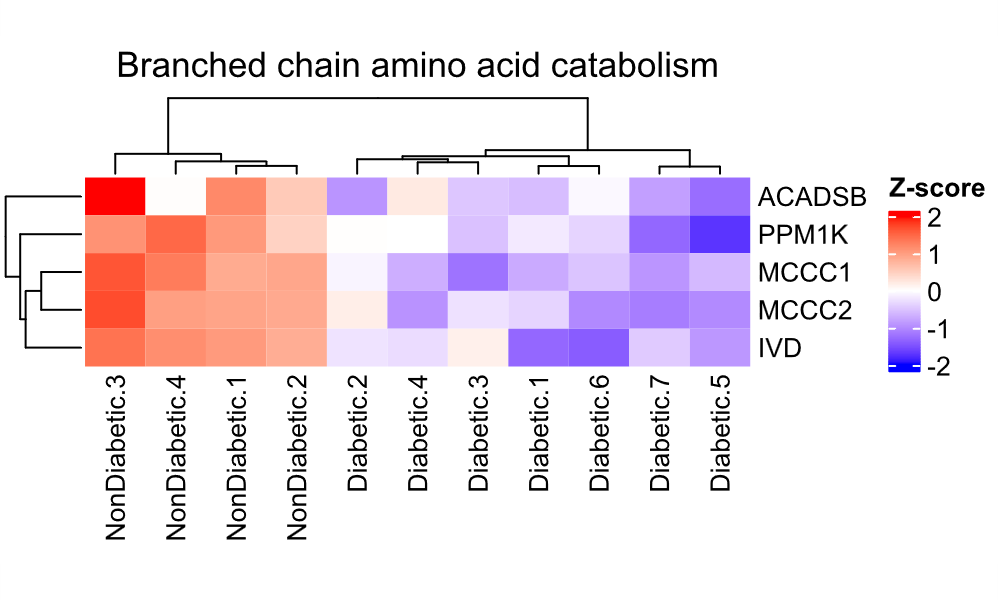

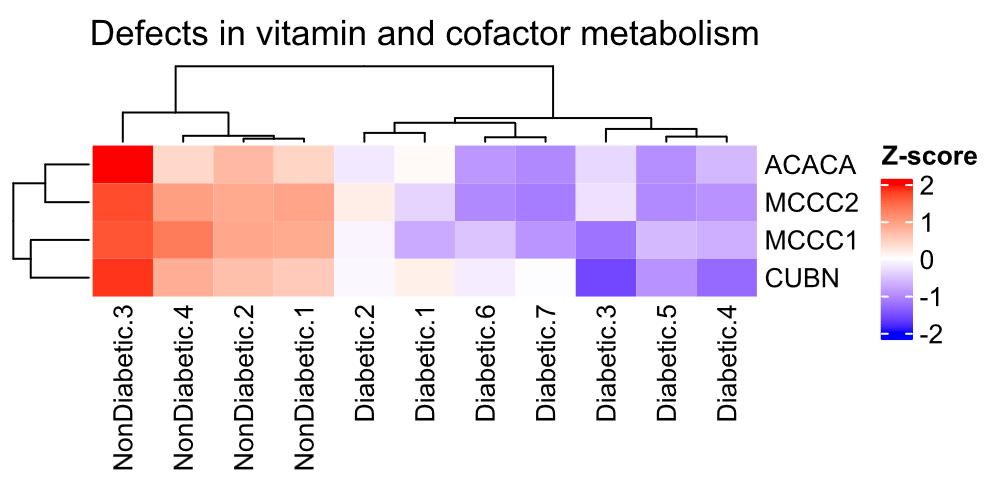

**A**

**B**

**C**

**D**

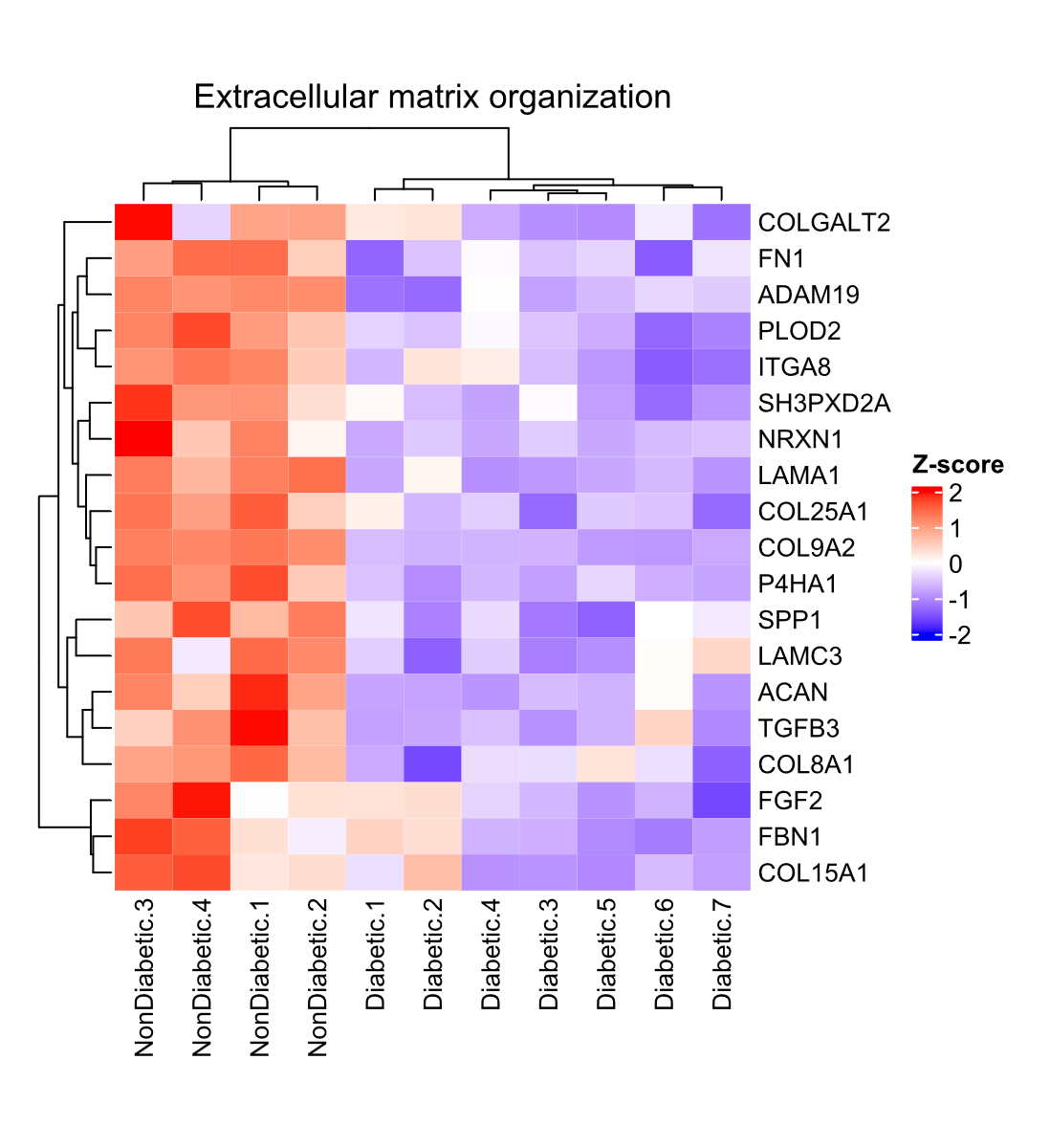
Figure S16. Downregulated Reactome enrichment – Clustered heatmap of genes downregulated in diabetic relative to nondiabetic wounds within the extracellular matrix organization pathway.

1	Velander, P. *et al.* Impaired wound healing in an acute diabetic pig model and the effects of local hyperglycemia. *Wound Repair Regen* **16**, 288-293 (2008). <https://doi.org:10.1111/j.1524-475X.2008.00367.x>

2	Maikawa, C. L. *et al.* A co-formulation of supramolecularly stabilized insulin and pramlintide enhances mealtime glucagon suppression in diabetic pigs. *Nature Biomedical Engineering* **4**, 507-517 (2020). <https://doi.org:10.1038/s41551-020-0555-4>

3	Christoffersen, B. O. *et al.* Functional and morphological renal changes in a Gottingen Minipig model of obesity-related and diabetic nephropathy. *Sci Rep* **13**, 6017 (2023). <https://doi.org:10.1038/s41598-023-32674-6>

4	Gerrity, R. G., Natarajan, R., Nadler, J. L. & Kimsey, T. Diabetes-Induced Accelerated Atherosclerosis in Swine. *Diabetes* **50**, 1654-1665 (2001). <https://doi.org:10.2337/diabetes.50.7.1654>

5	Wu, T. *et al.* clusterProfiler 4.0: A universal enrichment tool for interpreting omics data. *Innovation (Camb)* **2**, 100141 (2021). <https://doi.org:10.1016/j.xinn.2021.100141>

6	Patil, P. *et al.* Reactive oxygen species-degradable polythioketal urethane foam dressings to promote porcine skin wound repair. *Sci Transl Med* **14** (2022). <https://doi.org:10.1126/scitranslmed.abm6586>
